# 3D Reconstruction of Dinoflagellate Chromosomes from Hi-C Data Challenges the Cholesteric Liquid Crystal Hypothesis

**DOI:** 10.1101/2025.01.24.634729

**Authors:** Lucas Philipp, Georgi K. Marinov, Stephen Todd, Stephanie C. Weber

## Abstract

Dinoflagellates have permanently condensed chromosomes that are often described as liquid crystalline. Specifically, a Cholesteric Liquid Crystal (CLC) model was proposed in which DNA is organized into parallel fibers within stacked discs, such that the fiber orientation rotates by a constant angle between adjacent discs. Extrachromosomal loops extending from the discs were hypothesized to be more accessible and thus to contain transcriptionally active genes. Although the CLC model captures some features of dinoflagellate chromosome structure, its validity has not been rigorously tested against modern genomic data. Here, we use chromatin conformation capture (Hi-C) data to simulate 3D conformations of chromosome scaffolds for three dinoflagellate species: *Fugacium kawagutii*, *Symbiodinium microadriaticum*, and *Breviolum minutum*. Consensus and population-based modeling generate diverse polymer conformations with moderate orientational and nematic order. However, we find no evidence of cholesteric discs. Moreover, contact probability curves from empirical Hi-C data are inconsistent with the CLC model. Nevertheless, we show that introducing locus-specific boundaries into the CLC model can produce simulated Hi-C contact maps with topologically associating domains (TADs), which are observed in experimental Hi-C contact maps for these species. Finally, by mapping RNA-seq data onto our simulated conformations, we show that actively transcribed genes are present throughout entire chromosomes, and not exclusively on extrachromosomal loops or at the surface. Our results challenge the long-standing CLC model and suggest that dinoflagellate chromosomes are organized into condensed but non-crystalline structures that do not impede transcription.

## Introduction

Dinoflagellates are a diverse group of unicellular algae whose chromosomes remain highly condensed throughout the cell cycle [1]. How this unusual chromosome structure impacts gene expression is a long-standing question, since compact chromatin domains typically exclude RNA polymerase [2, 3] and suppress transcription [4] in other eukaryotes. The leading hypothesis to explain both the unique structure of dinoflagellate chromosomes and its consequences for transcription is the cholesteric liquid crystal (CLC) model. Originally proposed over fifty years ago, the CLC model posits that DNA is packaged into a series of flat discs that stack to form the long axis of the chromosome [5]. Within each disc, chromatin fibers are arranged parallel to a director that rotates by a constant angle between discs [6–9]. The CLC geometry is consistent with banding patterns observed in 2D transmission electron micrographs [5, 6]. In addition, extrachromosomal loops, described previously as 3D protrusions from the chromosome core, were hypothesized to connect both adjacent and non-adjacent discs and to contain transcriptionally-active genes, due to their presumed greater accessibility compared to the condensed chromosome core [5–8, 10–15]. However, the validity of the CLC model and its predictions for the spatial organization of transcription have not been rigorously tested against modern genomic data.

Chromosome conformation capture (Hi-C) is an experimental sequencing technique that measures which regions of primary DNA sequence contact each other in three-dimensional space. Hi-C analysis of several dinoflagellate species [16–18] revealed topologically associating domains (TADs), regions of primary sequence that interact with each other at much higher probabilities than with other chromosomal regions. These “dinoTADs” correlate with another common feature of dinoflagellate genomes, tandem arrays of co-oriented genes [19–22]. Indeed, the insulation boundaries that separate dinoTADs occur at locations where two oppositely-oriented gene arrays converge [16, 17]. Moreover, these boundaries are disrupted by transcription inhibitors, suggesting that active transcription may contribute to dinoTAD formation. Nevertheless, it remains unclear whether dinoTADs correspond to cholesteric discs and whether the CLC model is consistent with Hi-C data more generally [23].

Various computational methods have been developed to reconstruct 3D polymer conformations from Hi-C data [24, 25]. In the consensus modelling approach, all Hi-C-derived constraints are applied to a single conformation and contact probability is inversely related to spatial distance between a particular pair of monomers [25–29]. By contrast, in population-based modelling, contacts are distributed across many individual conformations in a population [30–32]. In this case, contact probability corresponds to the fraction of conformations in which a particular monomer pair is constrained to be in close proximity. Consensus modelling is fast and efficient, and therefore suitable for surveying hundreds of genomic regions across multiple species. Population-based modelling is computationally more expensive, but explicitly considers conformational heterogeneity which may result in polymer conformations that more closely reflect those found in individual cells [33, 34].

In this article, we use previously published Hi-C data [16–18] and both consensus [26] and population-based [32] modeling tools to simulate 3D conformations of chromosome scaffolds from three dinoflagellate species: *Fugacium kawagutii* (formerly known as *Symbiodinium kawagutii* [35]), *Symbiodinium microadriaticum* and *Breviolum minutum*. Surprisingly, our results indicate that dinoflagellate chromosomes are not organized into a stack of cholesteric discs as predicted by the CLC model. Rather, they contain multiple compact domains, and exhibit more orientational and nematic order than expected by chance, but less than CLCs. By mapping RNA-seq reads [36–38] onto our simulated polymer conformations, we find that active genes are located throughout chromosomes and not solely at their surfaces. This distribution suggests that although dinoflagellate chromosomes are highly condensed, they remain accessible to RNA polymerase. Our findings clarify important aspects of dinoflagellate chromosome structure and resolve a long-standing question as to how this structure relates to transcriptional activity.

## Results

### The CLC model does not produce TADs

The CLC model consists of a stack of cholesteric discs (Fig. 1 A) [5–9]. Within each disc, the chromatin fiber weaves back and forth in a parallel fashion. The director, along which the fiber is oriented, rotates by a constant angle, *θ*, between adjacent discs, thereby defining the cholesteric pitch. To compare the CLC model to empirical Hi-C data [16–18], we generated a population of 10^6^ CLC structures to approximate the number of cells sequenced in a typical bulk Hi-C experiment [39] (Methods). We assumed that discs were arranged consecutively in the stack, in the same order as in primary sequence (Fig. 1 A). To introduce conformational heterogeneity, we randomly incorporated extrachromosomal loops that extend outward from the stack of discs and re-enter either the same disc (intra-disc loops) or an adjacent disc (inter-disc loops) (Fig. 1 B). The length of each extrachromosomal loop was drawn from an exponential distribution. The variable positions and lengths of extrachromosomal loops resulted in variable locations of cholesteric discs along the primary sequence of individual CLC structures in the population (Supplementary Fig. S1 A).

**Figure 1.**
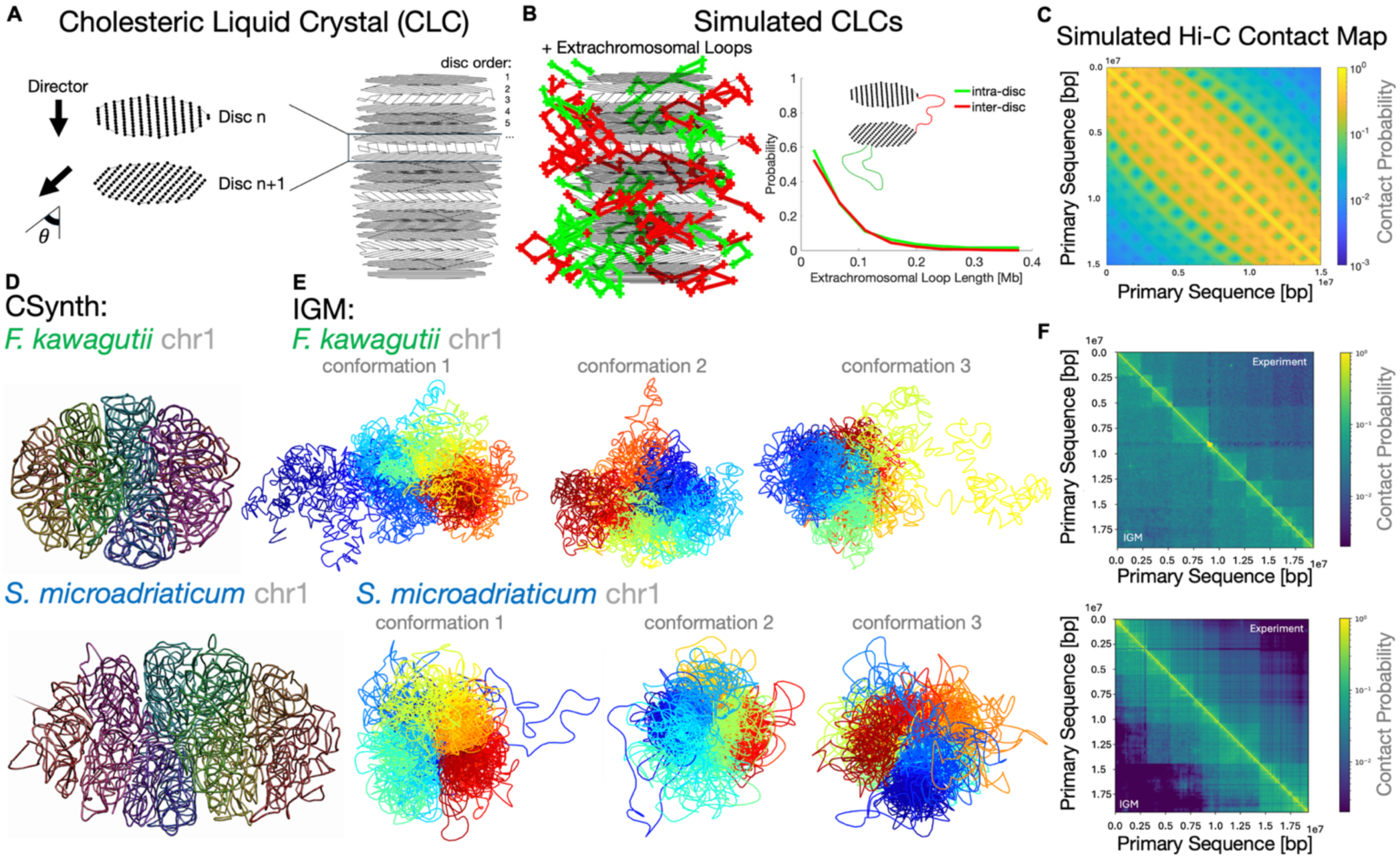
Dinoflagellate chromosomes lack cholesteric discs. A) The Cholesteric Liquid Crystal (CLC) model. The polymer aligns with a director (black arrows), which rotates by a constant angle 𝜃 between cholesteric discs. Discs are arranged in a consecutive order. B) CLCs are simulated by adding extrachromosomal loops to the stack of discs. Loops are positioned randomly and have exponentially distributed lengths. Intra-disc loops start and end within the same disc (green) and inter-disc loops connect vertically adjacent discs (red). C) Simulated Hi-C contact map for a population of 10^1^simulated CLC structures. D) CSynth conformations at 5 kb per monomer resolution of chromosome scaffold 1 from *Fugacium kawagutii* (Hi-C data: [18, 88]) and *Symbiodinium microadriaticum* (Hi-C data: [16]). Color indicates primary sequence. For the complete set of consensus conformations of dinoflagellate chromosome scaffolds, see Supplementary Figs. S4, S5. E) IGM conformations at 5 kb per monomer resolution from a population of 100 total conformations of the same chromosomes as in D. For a complete set of population-based conformations, see Supplementary Figs. S6, S7. F) Empirical Hi-C contact maps of representative dinoflagellate chromosomes (data: [16, 18, 88], upper right triangles) and simulated Hi-C contact maps from IGM populations (lower left triangles). Pearson correlation coefficients: *r* = 0.91 for *F. kawagutii*, *r* = 0.84 for *S. microadriaticum*.

We simulated a Hi-C contact map from our population of CLC structures and found a grid-like pattern, with stripes of high contact probability that are parallel to each other and perpendicular to the central diagonal (Fig. 1 C). The number and spacing of stripes are determined by the number and diameter of cholesteric discs in the input structures (Supplementary Fig. S1 B, S2 A, B). The grid-like pattern is robust to changes in CLC model parameters, such as the percentage of total sequence in extrachromosomal loops, the cholesteric pitch, and the tapering of disc diameter at the top and bottom of the stack (Supplementary Fig. S2 C, D, E). However, variation of the number of cholesteric discs per structure within the same population results in a blurring of the stripes (Supplementary Fig. S2 F).

The simulated Hi-C contact maps from CLC populations (Fig. 1 C, Supplementary Fig. S2) are qualitatively different from empirical Hi-C contact maps from the dinoflagellates *F. kawagutii* and *S. microadriaticum* [16–18] (Fig. 1 F, top right triangles). The most striking difference is the lack of TADs. Indeed, quantitative TAD metrics such as the insulation score [40] and directionality index [31] smoothly oscillate with a constant period for CLCs, while they vary irregularly and abruptly for dinoflagellate chromosomes (Supplementary Fig. S3). Overall, Hi-C contact maps from the CLC model are only modestly predictive of experimental data, with Pearson correlation coefficients *r* = 0.62 for *F. kawagutii* and *r* = 0.59 *S. microadriaticum*. These results indicate that the CLC model does not produce TADs, which are the dominant structural feature of dinoflagellates chromosomes [16–18]. **Dinoflagellate chromosomes lack cholesteric discs**

Next, we computed the 3D conformation(s) of dinoflagellate chromosomes that best reproduce empirical Hi-C contact probabilities. To mitigate algorithm bias [42], we used both consensus and population-based modeling approaches. First, we applied the CSynth software package [26], which incorporates Hi-C contacts as spring-like forces between monomers that serve to constrain their positions and outputs a single consensus conformation (Methods). After validation (see SI Text), we used CSynth to simulate conformations for all *F. kawagutii* (Supplementary Fig. S4, Supplementary Table S1) and *S. microadriaticum* (Supplementary Fig. S5, Supplementary Table S2) chromosome scaffolds at 5 kb per monomer resolution. Second, we implemented the Integrative Genome Modeling (IGM) software package [32], which uses maximum likelihood estimation to generate a population of conformations that is statistically consistent with bulk Hi-C data. We used IGM to simulate a population of 100 conformations for the first (largest) chromosome scaffold from *F. kawagutii* (Supplementary Fig. S6) and *S. microadriaticum* (Supplementary Fig. S7), also at 5 kb per monomer resolution.

In contrast to the CLC model (Fig. 1 A, B), we found that 3D conformations of dinoflagellate chromosomes lack cholesteric discs (Fig. 1 D, E). Instead, consensus conformations are compact and ellipsoidal (Fig. 1 D, Supplementary Figs. S4, S5), while population-based conformations can be elongated or spherical, with multiple compact domains and long extrachromosomal loops (Fig. 1 E, Supplementary Figs. S6, S7). We observed few extrachromosomal loops in consensus conformations – 0.63 per chromosome in *F. kawagutii* (Supplementary Fig. S4) and 1.8 per chromosome in *S. microadriaticum* (Supplementary Fig. S5) – but many more in the population-based conformations – 2.5 per conformation in *F. kawagutii* (Supplementary Fig. S6) and 2.9 per conformation in *S. microadriaticum* (Supplementary Fig. S7). Extrachromosomal loops were previously reported to be 30-60 kb in length [43], corresponding to 6-12 monomers per loop in our simulations. Therefore, the apparent sparseness of extrachromosomal loops in our consensus conformations is not due to limited resolution.

We next simulated Hi-C contact maps from our population-based (IGM) conformations of chromosome 1 for both species and found squares along the central diagonal, rather than a grid (Fig. 1 F, bottom left triangles). This pattern is consistent with dinoTADs [16–18]. Moreover, quantitative comparison of these contact maps with the experimental input data showed strong correlation (*r* = 0.91 for *F. kawagutii*, *r =* 0.84 for *S. microadriaticum*) (Fig. 1 F).

Finally, we simulated chromosome conformations for a third dinoflagellate species, *B. minutum*. While Hi-C data are available [17], they only cover an estimated 39% of the genome, compared to 79% for *F. kawagutii* and 64% for *S. microadriaticum* (Supplementary Table S3). Since *B. minutum* scaffolds may not correspond to full chromosomes, we selected the 10 largest Hi-C scaffolds for reconstruction (Supplementary Table S4). Consensus modeling generated spherical conformations with numerous extrachromosomal loops (5.3 loops per scaffold) (Supplementary Fig. S8 A), in contrast to the ellipsoidal conformations with few loops seen for *F. kawagutii* and *S. microadriaticum* (Fig. 1 D, Supplementary Fig. S4, S5). These conformational differences between species may be due to incomplete scaffolds, rather than true structural differences in *B. minutum* chromosomes.

Interestingly, in this species, population-based modeling generated similar conformations, with spherical shapes and many extrachromosomal loops (3.7 loops per conformation) (Supplementary Fig. S8 B). As for the other two species, we found strong correlation (*r* = 0.93) between Hi-C contact maps simulated from our IGM population and observed for *B. minutum* [17] (Supplementary Fig. S8 C), which both exhibit TAD-like squares rather than CLC-like stripes. Overall, our polymer simulations suggest that dinoflagellate chromosomes are not organized into stacks of cholesteric discs.

### Locus-specific boundaries can recapitulate TADs in CLCs

In mammalian cells, TADs arise from loop extrusion — which leads to conformational heterogeneity among cells (and over time) — with locus-specific boundaries [34, 44–46]. To determine whether the CLC model is compatible with TADs, we grouped cholesteric discs according to observed dinoTAD sizes [16] and introduced locus-specific boundaries in between groups. The order of discs was randomly permuted within each group but not between groups (Fig. 2 A). The simulated Hi-C contact map for CLCs with disc order permuted within groups shows squares of high contact probability along the central diagonal (Fig. 2 B), which are characteristic of TADs. Indeed, the insulation score and directionality index vary more irregularly and abruptly than for CLCs with consecutive disc order (Supplementary Fig. S3). Therefore, the presence of locus-specific boundaries, together with conformational heterogeneity across the population, can recapitulate TADs in CLCs.

**Figure 2.**
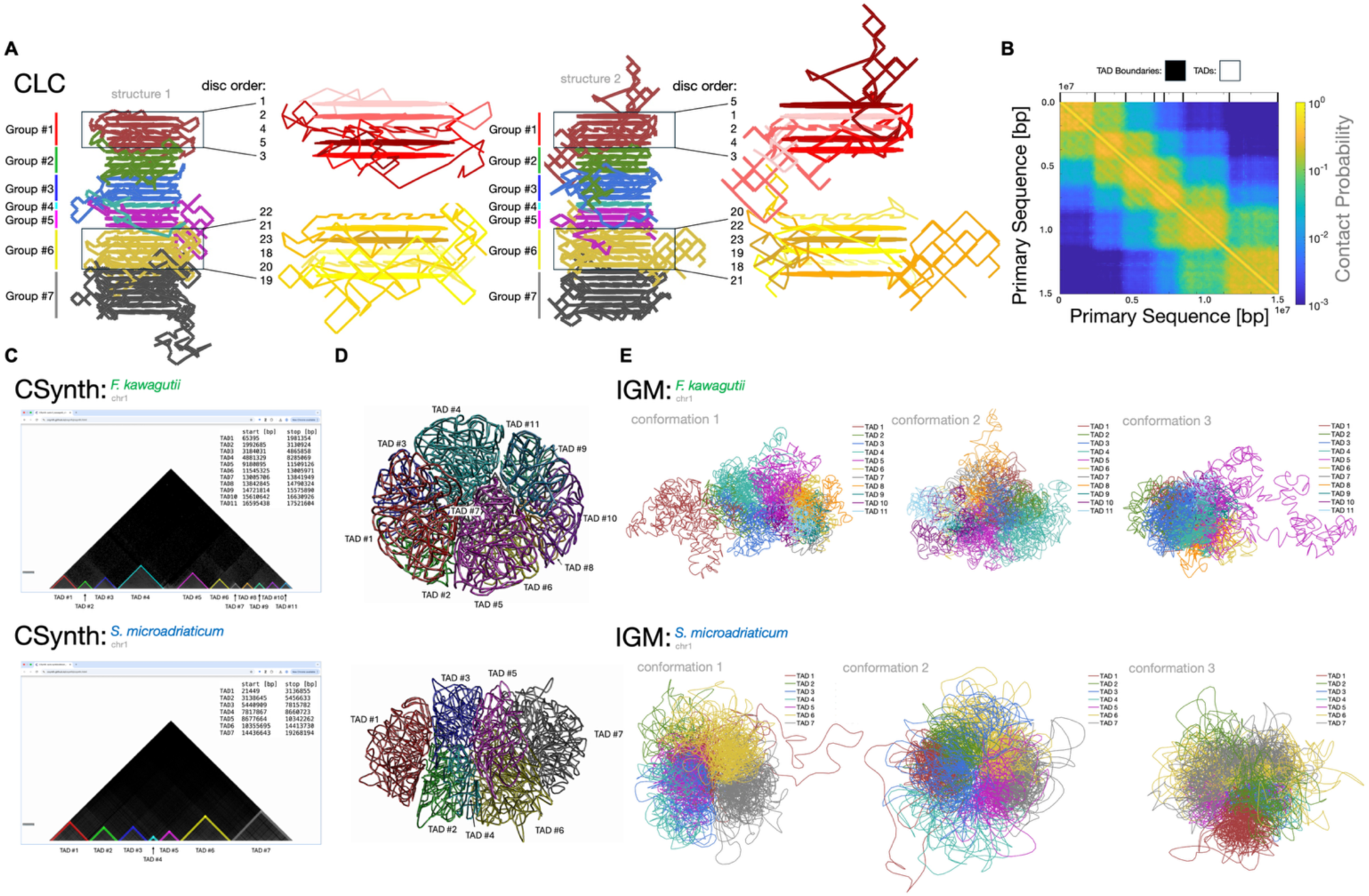
Locus-specific boundaries can recapitulate TADs in CLCs. A) Discs are grouped according to observed TAD size [16]. Insulating boundaries are added in between each group. Within each group, the primary sequence order of discs is permuted in the sub-stack across structures in the population. B) Simulated Hi-C contact map for CLCs with disc order permuted within groups. C) Manual annotation of TADs in CSynth generates a text file of boundary coordinates (inset). TADs can be visualized as distinct colors on the Hi-C contact map (bottom triangle) and in the 3D polymer conformation. D) TADs visualized using CSynth for *F. kawagutii* chr1 (top) and *S. microadriaticum* chr1 (bottom) at 5 kb per monomer resolution. E) TADs visualized in IGM conformations (3 of 100 total) of the same chromosomes.

Nevertheless, the correlation with the experimental *S. microadriaticum* contact map is only slightly better for CLCs with groups of permuted disc order (*r* = 0.67) than for CLCs with consecutive discs (*r* = 0.59, Fig. 1 C, F).

To visualize dinoTADs in our simulated chromosome conformations, we developed a manual TAD annotation tool within the CSynth graphical user interface. This tool allows the user to select the apex of a TAD with a mouse click and then automatically writes the left and right TAD boundary base-pair coordinates to a text file (Fig. 2 C, Methods). As with the rest of CSynth, this tool runs in a web browser and does not require any software installation [26]. Moreover, it can be used to annotate TADs in hundreds of chromosomes in less than a few hours. These TAD annotations can be used to visualize multiple TADs in distinct colors on the same 3D conformation (Fig. 2 D, Methods). Our manual TAD annotation tool has many advantages over existing automatic TAD annotators, which can be sensitive to parameter choice, involve lengthy computation times, require installation, or may not support integrated visualization of their output [47].

We used this new tool to annotate dinoTADs in Hi-C datasets [16–18]. A majority of the genome — 84% in *F. kawagutii* and 71% in *S. microadriaticum*, comparable to other eukaryotes [48, 49] — is organized into dinoTADs, which are located throughout our CSynth conformations but not in a regular arrangement or stack (Fig. 2 D). Similarly, dinoTADs in IGM conformations are not groups of cholesteric discs and exhibit diverse shapes depending on the conformation (Fig. 2 E). Therefore, although the CLC model can be made consistent with TADs, our 3D polymer conformations suggest that dinoTADs are not groups of cholesteric discs.

### Dinoflagellate contact probability curves are inconsistent with CLC model predictions

Contact probability curves are often used to describe chromosome organizational features [50]. This metric quantifies how frequently any two DNA loci, separated by a fixed amount of primary sequence, make contact in 3D space. Mathematically, contact probability can be written as 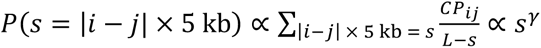 where 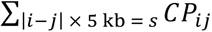 is the sum of Hi-C contact probabilities for all monomer pairs (𝑖, 𝑗) separated by 𝑠 = |𝑖 − 𝑗| × 5 kb, 𝐿 is the chromosome length, and 𝛾 is the contact probability decay exponent. The exact proportionality depends on the Hi-C normalization procedure. The decay exponent can be compared across species and to predictions from various models [50]. One such model is the equilibrium globule, constructed by first performing a 3D random walk within the confines of a sphere, and then turning on excluded volume interactions and thermally equilibrating the polymer. This procedure generates a knotted, space-filling polymer conformation [50]. Equilibrium globules have a decay exponent of 𝛾 = −1.5, while human interphase chromosomes have a decay exponent of 𝛾 = −1 [50].

To determine the decay exponent expected from the CLC model, we calculated the contact probability curve for a population of CLC structures following the methods outlined in [51]. For intermediate genomic separations, 𝑠 between 10^4^ and 10^6^ bp, 𝑃(𝑠) is very shallow with a decay exponent of 𝛾 ≈ −0.2 (Fig. 3 A). This exponent is robust across a range of CLC parameters (Supplementary Fig. S2), reflecting the high degree of compaction in this polymer model. At larger genomic separations, 𝑠 > 10^6^ bp, the curve decays more rapidly, consistent with the linear stacking of cholesteric discs [16]. Furthermore, contact probability curves for CLCs with consecutive disc order have two prominent local maxima at genomic separations corresponding to intra-disc (∼ 10^5^ bp) and inter-disc (∼ 10^6^ bp) contacts (vertical dashed lines: Fig. 3 A, Supplementary Fig. S2). CLCs with permuted disc order within groups exhibit the same very shallow decay exponent of 𝛾 ≈ −0.2 as CLCs with consecutive disc order (Fig. 3 A). However, the second peak corresponding to inter-disc contacts vanishes. This is expected as discs that are adjacent in primary sequence are no longer adjacent in 3D space and therefore do not regularly contact each other.

**Figure 3.**
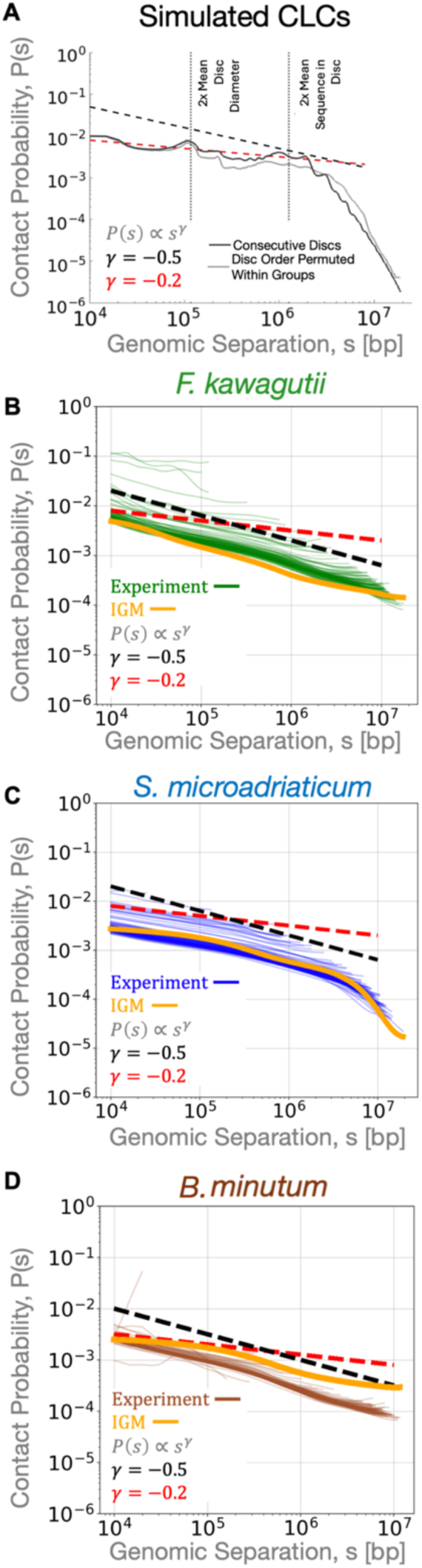
Contact probability curves for dinoflagellate chromosome scaffolds are inconsistent with CLC model predictions. A) Contact probability curve for a population of 10^1^ CLC structures with either consecutive disc order or disc order permuted within groups. The genomic separations corresponding to twice the mean cholesteric disc diameter and mean for two cholesteric discs are indicated by vertical dotted lines. Black dashed line represents a power law relationship with an exponent 𝛾 ≈ −0.5. Red dashed line represents a power law with exponent 𝛾 ≈ −0.2. Contact probability curves for the 75 largest chromosome scaffolds for B) *F. kawagutii* (Hi-C data: [18]); C) *S. microadriaticum* (Hi-C data: [16]); D) *Breviolum minutum* (Hi-C data: [17]). Each curve corresponds to an individual chromosome scaffold. Orange lines correspond to contact probability curves for a population of 100 IGM conformations for chromosome 1.

Next, we calculated contact probability curves for *F. kawagutii* and *B. minutum*, as was done previously for *S. microadriaticum* [16], using the cooltools package [52]. We selected the largest 75 chromosome scaffolds for each species (Supplementary Tables S1, S2, S4), as they likely represent completely assembled chromosomes, and excluded shorter scaffolds from our analysis. 𝑃(𝑠) curves for dinoflagellate chromosome scaffolds are smooth, with a decay exponent of 𝛾 ≈ −0.5 (Fig. 3 B-D). Contact probability curves from simulated IGM conformations produce the same 𝛾 ≈ −0.5 decay exponent (Fig. 3 B-D). This power-law scaling is steeper than what we observed for the CLC model (Fig. 3 A, Supplementary Fig. S2), even when groups of permuted discs produce TADs in the CLC contact map (Fig. 2 B), suggesting that dinoflagellate chromosomes are less condensed than CLCs.

Moreover, the experimental and IGM contact probability curves have no peaks at intermediate genomic separations, as would be expected for cholesteric discs. Overall, we find no evidence of cholesteric organization in the contact probability curves of dinoflagellate chromosome scaffolds. Although the CLC model can be made consistent with TADs, these discrepancies indicate that a stacked organization of cholesteric discs is fundamentally incompatible with dinoflagellate Hi-C data.

### Dinoflagellate chromosomes exhibit moderate orientational and nematic order

Despite their apparent lack of cholesteric discs, we next sought to quantify the degree of order in our simulated chromosome conformations. We first quantified the orientational order within our consensus and population-based conformations by calculating the orientational order parameter (OOP) [53] and its Fourier transform. The OOP can be defined as 𝑂𝑂𝑃(|𝑖 − 𝑗| × 5 kb) = < 𝑟⃗_’_ ⋅ 𝑟⃗_(_ >, where 𝑟⃗_’_ is the unit tangent vector to the 𝑖th monomer, and the brackets <⋅> denote the average dot product over all possible pairs of monomers (𝑖, 𝑗) separated by 𝑠 = |𝑖 − 𝑗| × 5 kb. For helical organization, the OOP is a sinusoid as a function of primary sequence separation and its Fourier transform is a narrow peak corresponding to the period of the helix (Fig. 4 A). For a particular genomic separation 𝑠, an OOP of 1 indicates perfect parallel alignment, an OOP of −1 indicates perfect anti-parallel alignment, and an OOP of 0 indicates random orientation or no alignment. Intermediate positive/negative OOP values indicate partial alignment/antialignment.

**Figure 4.**
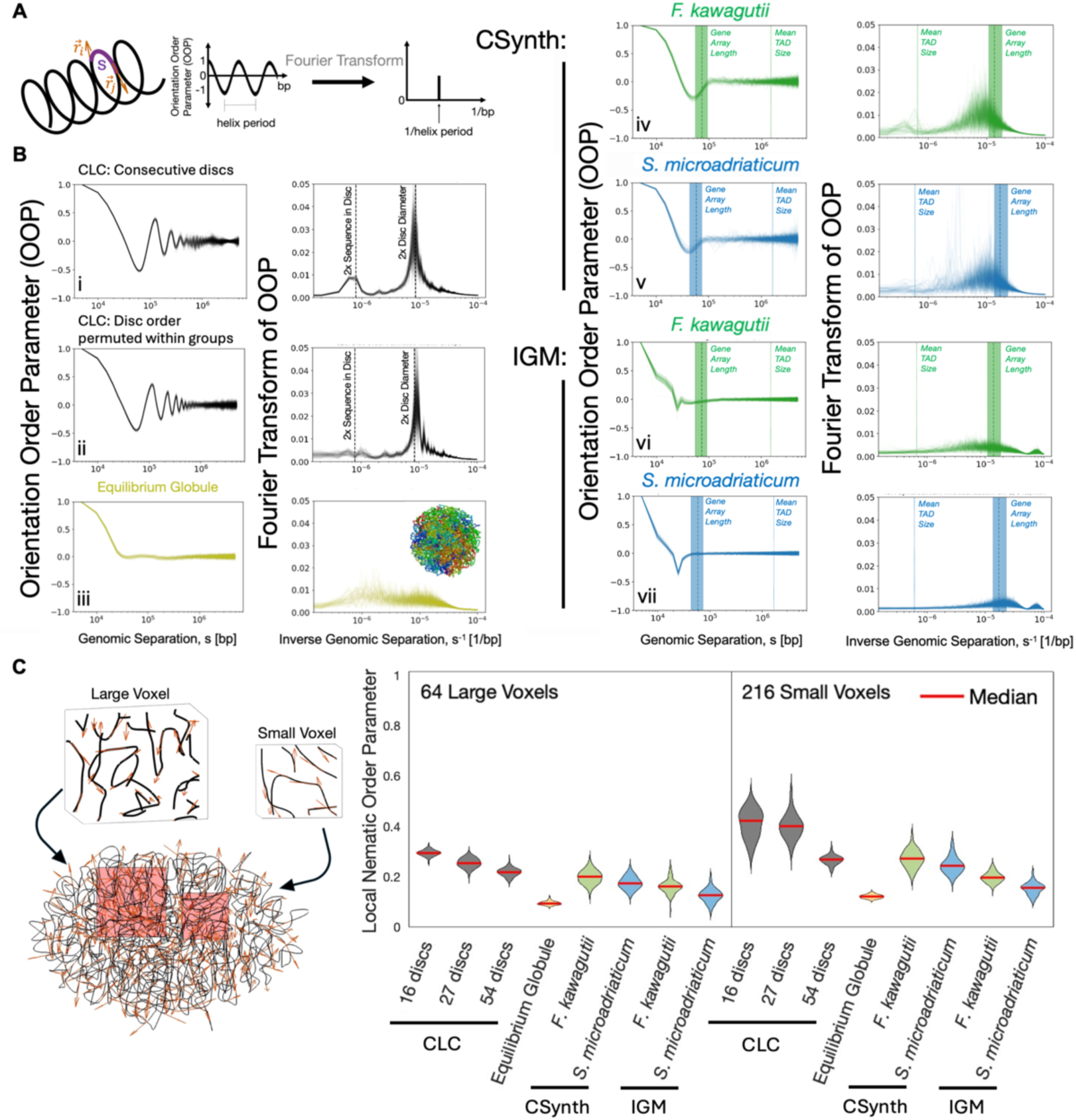
Dinoflagellate chromosomes exhibit moderate orientational and nematic order. A) Orientational order parameter (OOP) and its Fourier transform for a simple helix. Tangent vectors 𝑟⃗_i_ and 𝑟⃗_j_ (red arrows) at monomers *i* and *j*, located a primary sequence separation 𝑠 (purple) apart. B) Left: OOP. Right: Fourier transform of OOP. Each curve corresponds to one of 75 structures/conformations. Row i: CLCs with consecutive disc order. Row ii: CLCs with disc order permuted within groups. Dashed lines: reciprocal of the average amount of sequence in two cholesteric discs, reciprocal of the amount of sequence in twice the average disc diameter. Row iii: Equilibrium globules (data: [50]). Row iv: *F. kawagutii*, CSynth. Row v: *S. microadriaticum*, CSynth. Each curve corresponds to a consensus conformation of one of the 75 largest chromosomes. Row vi: *F. kawagutii*, IGM. Row vii: *S. microadriaticum*, IGM. Each curve corresponds to one conformation from a population of 100 total conformations of chromosome 1. The mean gene array size (±1 standard deviation) and the mean TAD size are indicated. C) Left: Example CSynth conformation of a dinoflagellate chromosome (black) with tangent vectors (red arrows). Small (edge length = 16.7% chr long axis) and large (edge length = 25% chr long axis) voxels (red boxes) are blown-up as insets and used to estimate short and long range orientational order, respectively. Right: Local nematic order parameter is calculated for all tangent vectors within every voxel and across 75 structures/conformations.

We compared the OOPs of CLCs as a positive control; equilibrium globules as a negative control; and simulated conformations of dinoflagellate chromosome scaffolds (Fig. 4 B, Supplementary Fig. 8 D). The OOP of CLCs with consecutive disc order exhibits a damped oscillation (Fig. 4 B, Row i). Two peaks are present in its Fourier transform: one peak corresponds to orientational ordering between adjacent cholesteric discs (∼10^−6^/bp), and the other corresponds to orientational ordering within a cholesteric disc (∼ 10^−5^/bp). The peak locations depend on the average number of monomers in a cholesteric disc, and not on the percentage of sequence in extrachromosomal loops, as expected (Supplementary Fig. S9). The OOP of CLCs with permuted disc order within groups also has a damped oscillation (Fig. 4 B, Row ii) but lacks the peak corresponding to inter-disc orientational alignment. This is expected because permutation of disc order results in discs that are adjacent in primary sequence but not in 3D space having directors that differ by random multiples of 𝜃 (Fig. 1 A).

Equilibrium globules are confined 3D random walks with no orientational order. We calculated the OOP of equilibrium globules using simulated data from [50]. As expected, the OOP monotonically decays at short genomic separations and plateaus to zero. There are no peaks in its Fourier transform, consistent with its lack of any periodic structure (Fig. 4 B, Row iii).

In contrast, the OOPs for consensus conformations of *F. kawagutii, S. microadriaticum* and *B. breviolum* chromosomes lack both the oscillatory signature of the CLC model and the monotonic decay of the equilibrium globule. Instead, they exhibit a negative peak at ∼ 10^4^ bp (Fig. 4 B, Rows iv & v, Supplementary Fig. S8 D), indicating partial anti-parallel orientation at this length scale. Since this corresponds to the average length of gene arrays, these peaks suggest that gene arrays may fold in an anti-parallel orientation in dinoflagellate chromosomes. Interestingly, we detect no orientational order on the scale of dinoTADs (∼10^6^ bp), likely due to their structural heterogeneity (Fig. 2 D, E). Fourier transforms of OOP curves for population-based conformations have smaller peaks at ∼ 10^−5^/bp (Fig. 4 B, Rows vi & vii, Supplementary Fig. S8 D), indicating less orientational order in IGM conformations compared to CSynth conformations.

Ordering of the chromosome could also occur independently of primary sequence separation and instead depend on the 3D spatial proximity of DNA loci. To measure this, we calculated the local nematic order parameter (NOP) at two different length scales following the methods outlined in [54]. A NOP of 1 indicates that all tangent vectors within a voxel are co-aligned in either a parallel or antiparallel orientation. A NOP of 0 indicates that tangent vectors within a voxel are randomly oriented with no directional correlation. Across all polymer conformations, the NOP decreases as voxel size increases because tangent vectors are averaged over a larger volume (Fig. 4 C). For CLCs, NOP also decreases as the number of discs increases because more discs with directors that are oriented differently are located within the same voxel (Fig. 4 C). For equilibrium globules, we found a negligible degree of nematic ordering, as expected (Fig. 4 C). Finally, we observe a moderate degree of nematic ordering in dinoflagellate chromosome conformations, intermediate between that of CLCs and equilibrium globules (Fig. 4 C, Supplementary Fig. S8 E). Altogether, both OOP and NOP metrics indicate that dinoflagellate chromosomes, simulated by either modeling approach, are more ordered than equilibrium globules, but less ordered than CLCs.

### Active genes occur throughout dinoflagellate chromosomes

Finally, we visualized the 3D spatial distribution of active genes in dinoflagellate chromosomes by mapping RNA-seq reads [36–38] to the genome assembly, and in turn, onto their corresponding 3D locations as predicted by CSynth and IGM. We defined a cylindrical coordinate system where the cylindrical axis is the chromosome’s long axis and the radial axis is all perpendicular directions from the cylindrical axis to the chromosome’s surface (Fig. 5 A). The CLC model — with or without locus-specific boundaries — is cylindrical in shape and predicts that active transcription occurs on extrachromosomal loops, which are located at the chromosome surface, as opposed to the core, which is hypothesized to be inaccessible to RNA polymerases [5–8, 10–15, 55–58]. Thus, we would expect to see the highest expression levels at the surface and low or no expression in the core (Fig. 5 A).

**Figure 5.**
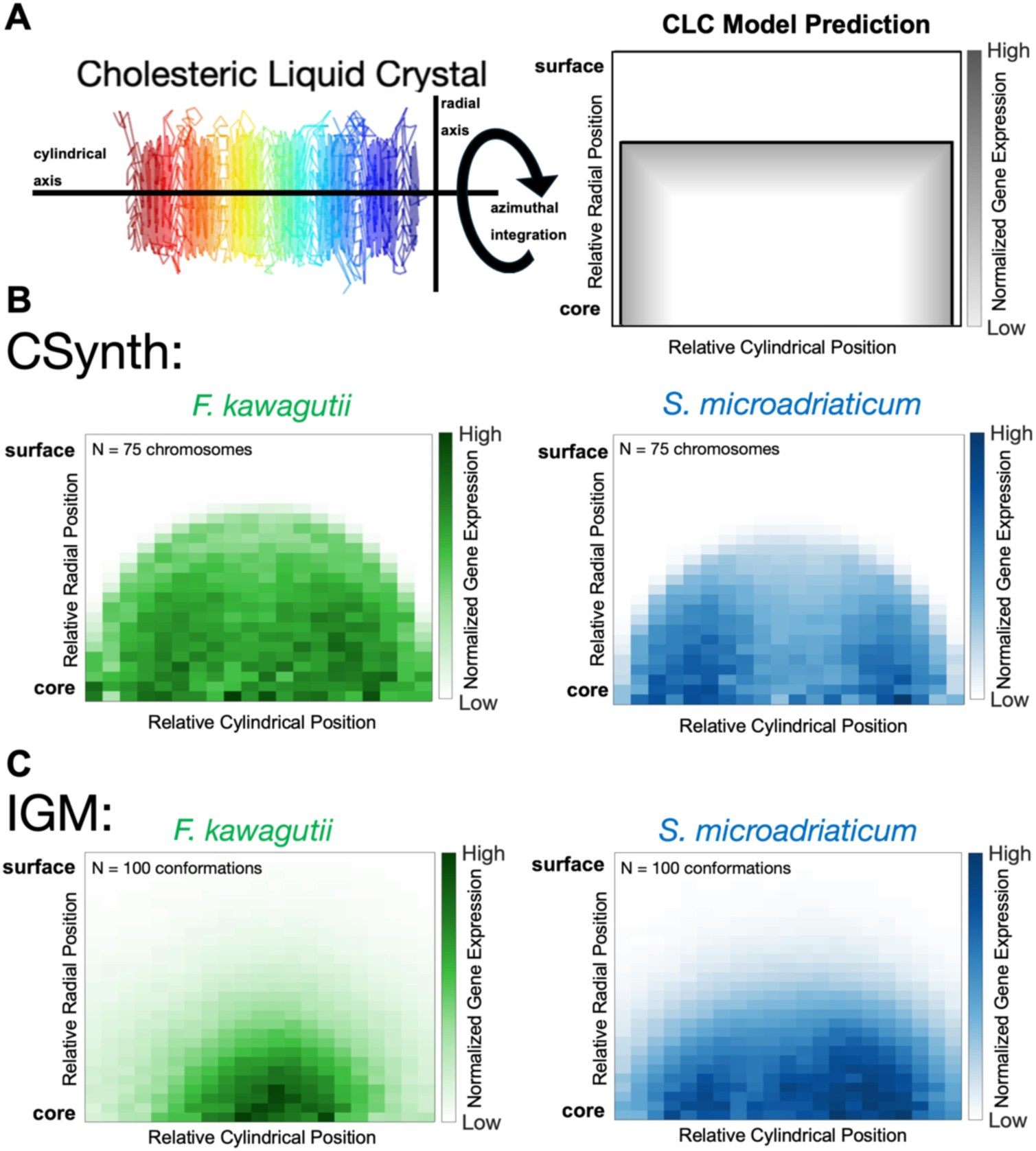
Active genes occur throughout dinoflagellate chromosomes. A) Cylindrical coordinate system with cylindrical axis aligned with the chromosome long axis. The outward radial axis is perpendicular to the cylindrical axis. The CLC model predicts that active genes are near the chromosome surface on extrachromosomal loops, and not in the condensed core. B) Normalized gene expression (data: [36–38]) as a function of distance to core (relative radial position) and position along long axis (relative cylindrical position) of consensus conformations for *F. kawagutti* (green) and *S. microadriaticum* (blue). C) Normalized gene expression of population-based conformations of chromosome 1 for each species using the same cylindrical coordinate system.

To determine whether transcription levels correlate with proximity to the surface, we used this coordinate system to calculate the average gene expression throughout *F. kawagutii* and *S. microadriaticum* chromosome conformations, controlling for variation in DNA density across voxels (Fig. 5 B-C, Methods). In contrast to the CLC model’s prediction, we observe active genes throughout dinoflagellate chromosomes modeled using both CSynth (Fig. 5 B) and IGM (Fig. 5 C). Transcription heatmaps of population-based conformations have more diffuse peripheral boundaries compared to consensus conformations, consistent with their more variable shapes and numerous extrachromosomal loops (Fig. 1 E, Supplementary Figs. S6, S7). Interestingly, we find somewhat increased transcription levels away from the center of the cylindrical axis for *S. microadriaticum* chromosomes (Fig. 5 B-C, blue). Telomeres are also preferentially located away from the center of the cylindrical axis (Supplementary Fig. S10), which is consistent with the previous observation that *S. microadriaticum* chromosomes have higher gene density at their telomeres [16]. Overall, we find no correlation between gene expression and radial position in dinoflagellate chromosomes, indicating that active transcription is not restricted to extrachromosomal loops.

## Discussion

We have systematically and quantitatively compared genomic data for three species of dinoflagellates to predictions of the CLC model. Unexpectedly, our results challenge the long-standing hypothesis that dinoflagellate chromosomes are organized into CLCs. First, empirical Hi-C contact maps have TADs [16–18] instead of a grid-like pattern that we show is characteristic of cholesteric discs (Fig. 1). Second, the scaling of contact probability curves for dinoflagellate chromosome scaffolds are inconsistent with CLCs (Fig. 3). Third, our polymer simulations predict 3D conformations that are less compact and less ordered than CLCs (Fig. 4). Finally, we find active genes located throughout these 3D conformations, rather than on the surface, as expected for CLCs (Fig. 5).

The CLC model was proposed to explain the unique structure of dinoflagellate chromosomes [5–9]. It successfully predicted a highly compact twisted chromosome, with significant nematic order, and became the accepted view [5–15, 64–66]. However, our systematic analysis of reconstructed chromosome conformations – as well as experimental data [16–18] – reveals several important disagreements between dinoflagellate chromosomes and the CLC model. Therefore, we conclude that dinoflagellate chromosomes (at least in these three species) are not cholesteric liquid crystals.

Different computational modeling approaches can generate different outputs from the same input data, creating uncertainties for interpretation [46]. To mitigate this risk, and to enhance the robustness of our conclusions, we used two different software packages to reconstruct dinoflagellate chromosomes. While both CSynth and IGM produce 3D polymer conformations that disagree with the CLC hypothesis, their predictions differ in some notable ways. For example, the ellipsoidal shape of CSynth conformations is reminiscent of dinoflagellate chromosomes observed by transmission electron microscopy [1, 5–8, 13, 59–63], while IGM conformations sometimes adopt dumbbells and other elongated or irregular shapes (Supplementary Figs. S6, S7) that are less common in the microscopy literature [1]. Nevertheless, IGM conformations give rise to simulated Hi-C contact maps (Fig. 1F) and contact probability curves (Fig. 3) that are highly consistent with empirical data [16–18]. Interestingly, CSynth can reproduce CLC-like conformations while IGM cannot (SI text, Supplementary Fig. S15). Overall, our work highlights both the strengths and limitations of these computational models. Finally, IGM is capable of simultaneously integrating Hi-C data together with data from orthogonal experimental methods including fluorescence in situ hybridization (FISH), DNA Adenine Methyltransferase Identification (Dam ID), and single-cell split-pool recognition of interactions by tag extension (SPRITE) [32]. While such data types are not yet available for dinoflagellates, optimizing these methods and applying them to these organisms should be a priority for future research.

The 3D conformations produced by CSynth and IGM are static. However, chromosomes are highly dynamic in living cells [46]. Analysis of temporal variation in chromosome structure requires Hi-C data from a synchronized population of cells sampled at different time points throughout the cell cycle [67, 68]. Comparison of Hi-C contact maps of *S. microadriaticum* cultures enriched for either flagellated mastigote cells (G1/S) or immobile coccoid cells (G2/M) showed no apparent differences, indicating that this species’ chromosomes are permanently condensed and undergo minimal rearrangements during the cell cycle [16]. Nevertheless, our study does not rule out the possibility of dynamic rearrangement of extrachromosomal loops to the chromosome exterior where they are hypothesized to be more transcriptionally accessible [69]. If this were the case, then the expression of nearly all genes would need to be regulated via this mechanism as nearly all chromosomal regions are transcriptionally active (Fig. 5 B, C). Alternatively, the dense core of dinoflagellate chromosomes may not exclude the transcriptional machinery after all. Indeed, recent Assay for Transposase-Accessible Chromatin using sequencing (ATAC-seq) data on *B. minutum* indicate that while overall accessibility is low, it is more uniform than in other eukaryotic genomes, consistent with the lack of nucleosome-based chromatin [70]. Tn5 transposase (53.3 kD) is much smaller than RNA Polymerase II (Pol II, 514 kD), so it is possible that Tn5 can access regions from which Pol II is excluded.

However, bacterial RNA polymerase (457 kD) is able to penetrate the highly condensed nucleoid in stationary-phase *E. coli* cells [71], which has also been described as liquid crystalline [72]. Together, these findings suggest that dinoflagellate chromosomes are broadly accessible, and could explain why active genes are located throughout the chromosome rather than preferentially at the surface (Fig. 5).

In addition to the CLC model, many other models for dinoflagellate chromosome organization have been proposed. One model consists of two independent helices coiled around each other [73]. Another model consists of a toroidal chromosome twisted into a double helix, with rounded hairpin loops at the chromosome ends rather than exposed telomeres [74]. A third model proposes that dinoflagellate chromosomes are polytene, i.e. they undergo many rounds of complete and/or partial replication without segregation, maintaining close alignment of sister chromatids [75]. A fourth model consists of a hierarchical helix with six degrees of hierarchy [76, 77]. A fifth model consists of a helical condensed core, with abundant extrachromosomal loops [78].

Our 3D conformations of dinoflagellate chromosomes are not consistent with any of these models. They do not consist of multiple interwoven helices, are not twisted toroids, are not helices of helices, and do not contain abundant extrachromosomal loops. Furthermore, dinoflagellate Hi-C contact maps do not contain an X-shape pattern that is characteristic of circular chromosomes [79], do not have variable contact probability along the main diagonal as is the case for chromosomes with polytene bands [80], and lack significant trans-chromosomal contacts that would be expected from interwoven helices involving multiple chromosomes [16–18].

In addition to dinoflagellates, condensed, twisted, and ordered interphase chromosome structures have been observed in at least three other biological contexts: mosquito chromosomes [81], cricket sperm chromosomes [54], and bacterial nucleoids [82]. The latter two examples are particularly interesting because, like dinoflagellates, they also lack nucleosomes. Indeed, dinoflagellate viral nuclear protein (DVNPs) and histone-like proteins (HLPs) replace histones [56, 57, 83] as the dominant DNA-binding protein in dinoflagellates. Histones are replaced by SNBPs (sperm-specific nuclear basic proteins) in sperm cells [54] and bacteria use a variety of nucleoid-associated proteins, including HU (heat unstable) proteins [84], to package their DNA. Together, these examples suggest that highly condensed, ordered chromosome structures may be more common in the absence of histones.

Beyond DNA structure, the CLC model also makes functional predictions, including that active genes are located on the chromosome surface in extrachromosomal loops and not in the condensed chromosome core [5–8, 10–15, 55–58]. These predictions are mainly based on radiolabeling studies, which show incorporation of ^3^H-adenine or ^3^H-uridine at the nucleolus [10, 11, 55], but not overlapping with condensed chromosomes. Our results, which leverage recent RNA-seq datasets, suggest that transcription is spatially ubiquitous. We find active genes throughout dinoflagellate chromosomes and not exclusively at the chromosome surface (Fig. 5 B, C). Our findings are consistent with other modern genomic assays such as ATAC-seq, which indicate that dinoflagellate genomes are uniformly accessible to transposases [70]. Moreover, transcription still occurs in highly condensed human mitotic chromosomes [4]. Thus, dinoflagellate chromosomes are likely accessible to RNA polymerase and condensation is not the primary mechanism for regulation of gene expression. Rather, dinoflagellate genes can be actively transcribed regardless of their chromosomal location.

Although our results indicate that chromosome conformation does not influence gene expression in dinoflagellates, transcription may contribute to chromosome organization. For example, we observe orientational order at length scales corresponding to gene arrays (Fig. 4 B). The negative peak in the OOP curve for dinoflagellate chromosomes is similar to that observed in model human chromosomes perturbed by extensible force dipoles that represent transcriptional activity [85]. Thus, transcription of co-oriented genes in divergent gene arrays may result in an anti-parallel arrangement. Transcription has also been implicated in dinoTAD formation. Indeed, the boundaries of dinoTADs in *S. microadriaticum* [16] and *B. minutum* [17] correspond to positions at which oppositely-oriented gene arrays converge and are sensitive to transcriptional inhibitors. We mapped RNA-seq reads [36–38] onto the *F. kawagutii* genome assembly and also found that dinoTAD boundaries occur between convergent gene arrays (Supplementary Fig. S11), consistent with these previous studies [16, 17].

Transmission electron microscopy (TEM) has been used frequently to generate 2D cross-sectional images of dinoflagellate chromosomes [1, 5–8, 13, 59–63]. To date, no study has imaged dinoflagellate chromatin fiber orientation in 3D. Instead, interpretation of the banding patterns observed in 2D cross-sectional images has relied on mathematically projecting 3D CLC models onto a 2D imaging plane [5]. The limitation of 2D data makes it difficult to extrapolate ordering observed in a single plane to volumes within a whole chromosome, and to track the contours of individual chromatin fibers as a function of primary sequence. For example, dinoflagellate chromosomes that produce banding patterns in TEM cross-sections, which are typically attributed to CLC organization, appear helical when viewed after a whole-mounting procedure [86]. In addition, a recent TEM study [62] found that while oblique and longitudinal sections of *Prorocentrum minimum* chromosomes agree qualitatively with CLC model predictions, chromatin fibers were not parallel within transverse sections — an observation that is inconsistent with cholesteric discs. The advantage of our Hi-C-based polymer modeling approach is that the entire dinoflagellate chromosome is modeled in 3D with the added context of primary sequence information.

Overall, this work advances our understanding of dinoflagellate chromosome organization in 3D, providing evidence that dinoflagellate chromosomes are not cholesteric liquid crystals. Future work should focus on 3D imaging and chromatin fiber tracing within dinoflagellate chromosomes, as well as 3D imaging of nascent RNA and transcription [87].

## Methods

### Hi-C data pre-processing

Hi-C-assisted genome assemblies and contact probabilities were accessed from the Gene Expression Omnibus (GEO) with accession numbers: GSE152150 [16] and GSE153950 [17], or from the Sequence Read Archive (SRA) with accession numbers: SRR25948349, SRR25948348 [88], and further processed following [18].

*F. kawagutii* Hi-C contact maps were converted from .hic format to .cool format using the hic2cool python package https://github.com/4dn-dcic/hic2cool.

When un-ligated and self-ligated Hi-C artifacts are abundant, Hi-C normalization can greatly reduce off-diagonal contact probabilities [89]. This results in weak CSynth Hi-C-based attraction forces and slow equilibration. Un-ligated and self-ligated Hi-C artifacts do not contain information about the 3D organization of the chromosome; they are short range contacts that map entirely within a single Hi-C bin (main diagonal), or across one Hi-C bin boundary (1st off-diagonal) [16]. To accelerate CSynth conformation equilibration, the ratio of average main diagonal contact probabilities 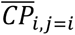 to average off-diagonal contact probabilities 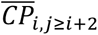 was set to 100 by dividing all main diagonal contact probabilities by 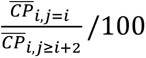, and the ratio of average 1st off-diagonal contact probabilities 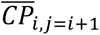 to average off-diagonal contact probabilities 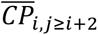 was set to 10 by dividing all 1st off-diagonal contact probabilities by 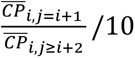. These modifications were chosen such that after normalization, CSynth Hi-C attraction forces remained high enough to allow conformation equilibration within minutes. Main diagonal and 1st off-diagonal contact probabilities were excluded for all contact probability curve calculations.

Individual Hi-C contact files for each chromosome were then normalized by Iterative Correction and Eigenvector decomposition (ICE) [89] and written to a text file in a CSynth compatible format via the cooler python package [90].

### Generating 3D chromosome conformations with CSynth

CSynth was accessed online at: https://csynth.github.io/csynth/csynth.html. The maximum number of monomers supported by our system’s GPU was determined by typing the command gl.getParameter(gl.MAX\_TEXTURE\_SIZE) into the console of the web browser’s developer tools. The following optimized parameters (SI Text) were used:

- Spring power: 0. This sets the relative importance of long and short distance effects.
- Contact force: 60. This sets the magnitude of attraction between Hi-C contacts.
- Push-apart power: −4. This sets the exponent for a global repulsive force between monomers that decays with distance. This exponent is analogous and has a similar value to the empirically derived exponent relating Hi-C contact probability and mean spatial distance determined by FISH in human IMR90 cells on chromosome 21 [91, 92].

After equilibration, the xyz coordinates of all monomers were saved to a text file. A picture of the equilibrated structure was saved separately.

### Simulating CLC structures

3D polymer models of CLCs were generated using a custom MATLAB script. Polymer length (20 Mb, 4000 monomers) was chosen to match the largest *S. microadriaticum* and *F. kawagutii* Hi-C scaffolds. Cholesteric discs were made by placing monomers on a square grid inside a circle of a chosen diameter. Monomers were connected in primary sequence by “weaving” the polymer backbone back and forth, parallel to a rotating director. The number and length of extrachromosomal loops were randomized while maintaining the total amount of sequence in loops at a user-defined fraction. After specifying the fraction of total monomers in extrachromosomal loops, and the fraction of loop monomers in inter-disc loops (1 minus the fraction of loop monomers in intra-disc loops), the remaining monomers were divided by the number of cholesteric discs minus 1 to give the mean inter-disc loop length. Inter-disc loop lengths were then drawn from an exponential distribution with a parameter determined by the mean inter-disc loop length. This was done to make loop starts/stops as uniformly distributed in monomer space as possible. Intra-disc extrachromosomal loop lengths were drawn from an analogous exponential distribution.

The primary sequence order of discs was either consecutive from top to bottom of the stack or permuted within groups but not across locus-specific boundaries which separate groups adjacent in primary sequence. In the case of CLCs with permuted disc order, inter-disc extrachromosomal loops were assigned to connect disc pairs according to that disc pair’s spatial distance separation rank because adjacent discs in primary sequence located further away in 3D space require longer loops to connect them.

### Simulating Hi-C contact maps

For CLCs and Csynth, Hi-C contacts were simulated using 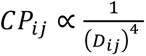 [91, 92] where 𝐶𝑃*_ij_* represents the Hi-C contact probability for genomic loci pair (𝑖, 𝑗), and 𝐷_’(_ represents the Euclidean distance between these two monomers. For population level Hi-C contact maps, the contact probability for each single-conformation Hi-C contact map was averaged per pixel (𝑖, 𝑗) across all conformations.

For IGM, Hi-C contact maps were simulated as previously described in [32].

### Quantitative TAD metrics

We used the FanC python package [93] to calculate insulation score [40] and directionality index [41] using a Hi-C bin size of 5 kb and a window size of 100 kb.

### Equilibrium globule structures

Equilibrium globule structures were downloaded from GEO using the accession number: GSE18199 [50].

### Analyzing spatial organization of transcription

RNA-seq data from asynchronous cell populations were accessed via the SRA using the following accession numbers: for *S. microadriaticum*, SRR3337493 [36]; for *F. kawagutii*, SRR9417753, SRR9417755, SRR9417756, SRR1300302, SRR1300303, SRR1300304, and SRR1300305 [37, 38].

Alignment to the genome assembly was performed following the methods outlined in [17].

### Calculating normalized gene expression

The largest 75 chromosome scaffolds, likely representing complete or near-complete chromosome assemblies, for each dinoflagellate species were scaled isotropically to the same size, with each chromosome long axis aligned horizontally. Using a cylindrical coordinate system, the total number of RNA-seq reads were counted in each voxel, integrated azimuthally, and divided by the cylindrical Jacobian which controls for variation in amount of DNA per voxel.

### Annotating TADs manually with CSynth

After loading a Hi-C contact file into CSynth, the chromosome was hidden and its dynamics were arrested. Hi-C contacts were displayed on CSynth’s matrix viewer. The *matdistfar* parameter was adjusted until TADs were clearly visible and rotation was set to 1.5 for a more upright matrix view. To annotate a TAD, the TAD apex was selected using the mouse, and edges then appeared outlining the TAD boundary. TAD coordinates were saved separately for each TAD using option+Z (Macintosh) or alt+Z (Windows), and collectively exported in BED file format using shift+option+Z (Macintosh) or shift+alt+Z (Windows). Finally, TADs were colored in 3D after un-hiding the chromosome and dragging the .bed file into the CSynth browser window. For a detailed list of commands, see SI Text.

## Data availability

Code is available at: https://github.com/lucasphilipp1/3D-Reconstruction-of-Dinoflagellate-Chromosomes-from-Hi-C-Data. *F. kawagutii* Hi-C data, *F. kawagutii* and *S. microadriaticum* CSynth conformations, CLC structures, RNA-seq .bed files, and TAD .bed files, have been deposited to https://doi.org/10.5281/zenodo.14285613.

## Supplementary Information Appendix (SI)

Supplementary Information accompanying this article including SI Text, Supplementary Figures S1-S15 and Tables S1-S4 can be found here.

## Acknowledgements

We would like to thank Audrey Baguette for helpful discussions regarding Hi-C data pre-processing. L.P. was supported by a Canadian Graduate Scholarship – Master’s and a Canadian Graduate Scholarship – Doctoral from the National Science and Engineering Research Council of Canada. The work was funded by the New Frontiers in Research Fund (NFRFE-2019-00189 to S.C.W.). This research was undertaken, in part, thanks to funding from the Canada Research Chairs Program.

## Author contributions

L.P. and S.C.W. designed the research, wrote the paper, and obtained funding; G.K.M. improved the *F. kawagutii* genome assembly using raw Hi-C reads, assembled *F. kawagutii* contact maps, and aligned RNA-seq reads to the genome; S.T. added data visualization and manual TAD annotation functionalities to CSynth; L.P. performed all other research aspects.

