## Supplementary Information for "3D Reconstruction of Dinoflagellate Chromosomes from Hi-C Data Challenges the Cholesteric Liquid Crystal Hypothesis"

- SI Text
- Figures S1 to S15
- Tables S1 to S4
- SI References

SI Text:

**Model Validation**

We validated CSynth in three different ways. First, we generated a CLC structure with known monomer positions to serve as a positive control (Supplementary Fig. S12 A). We calculated a Hi-C contact map from this CLC structure (Supplementary Fig. S12 B) and fed it into CSynth. The CSynth-generated output conformation (Supplementary Fig. S12 C) consists of a rotating stack of ordered discs and has a 7% relative error in monomer pairwise separation compared to the known input CLC structure.

Second, we used empirical Hi-C [S1] and fluorescence in situ hybridization (FISH) [S2] datasets for chromosome 21 in human IMR90 cells, which show a strong correlation between contact probability and mean spatial distance, to determine the optimal parameter set that minimizes the relative error between the pairwise separation predicted by CSynth and real distances measured by light microscopy (652 FISH probes spaced 50 kb apart) (Supplementary Fig. S13).

Third, we applied this parameter set to reconstruct a wide variety of chromosome architectures using Hi-C data from diverse species [S3]. CSynth successfully captured a range of previously reported architectures, including the telomere-to-centromere axis in yellow fever mosquito chromosome 1 [S4] (Supplementary Fig. S14 A), telomere clustering in peanut chromosome 2 (Supplementary Fig. S14 B), and the absence of Rabl-like features (defined as centromere or telomere clustering [S3]) in roundworm chromosome 1 (Supplementary Fig. S14 C), with relative errors of 10.6%, 8.3%, and 11.4%, respectively.

Finally, we simulated the Hi-C contact map from a population of 100 CLC structures and used it as input to CSynth and IGM (Supplementary Fig. S15). The consensus conformation generated by CSynth has a cylindrical shape, with global twist and local alignment of the polymer perpendicular to the long axis, that is reminiscent of the CLC model. The simulated Hi-C map from the CSynth output is strongly correlated with the input map ($r = 0.82$). In contrast, the population-based conformations generated by IGM, while cylindrical, have some local polymer alignment but it is parallel to the long axis, and there is no global twist or stacking. Moreover, the similarity of the input and output Hi-C contact maps is poor ($r = 0.49$). Thus, in the case of CLCs, which are highly condensed, consensus modeling performs better than population-based modeling in capturing structural features and reproducing Hi-C contact maps.

**Annotating TADs manually with CSynth**

1. Uncheck "ribbon" and "simulation" in the GUI's drop-down menu to hide the chromosome and arrest its dynamics respectively.
2. Display Hi-C contacts using the menu: matrix→color→input A: "name of Hi-C file".
3. Adjust the *matdistfar* parameter until TADs are clearly visible.
4. For a more upright matrix view, increase matrix→rotation to 1.5.
5. To annotate a TAD, select the TAD apex using the mouse and edges will appear outlining the TAD boundary. Then press option+Z (Macintosh) or alt+Z (Windows) to save the TAD coordinates and repeat for other TADs.
6. Press shift+option+Z (Macintosh) or shift+alt+Z (Windows) to download a .bed file with the TAD start/end basepair coordinates as rows.
7. To clear the memory of TAD coordinates, press and hold M, and then press zero.
8. To color TADs in 3D, after re-checking "ribbon", drag and drop the .bed file into the CSynth browser window.

**Calculating Relative Error**

Relative error was used to quantify the similarity between Hi-C contact maps and geometric similarity between chromosome conformations, as has been done previously [S5, S6].

To compare a known CLC structure to a CSynth conformation predicted from the CLC structure's Hi-C contact map (Supplementary Fig. S12), we calculated Relative Error = $\frac{2}{N(N+1)}\sum_{\mathrm{i,j \geq i}}^{\mathrm{N}} \frac{|\mathrm{Input\ separation_{ij}} - \mathrm{CSynth\ separation_{ij}}|}{\mathrm{Input\ separation_{ij}}} \times 100\%$ , where $\mathrm{Input\ separation_{ij}}$ [a.u.] is the Euclidian distance between monomers $(i, j)$ in the assumed CLC structure and $\mathrm{CSynth\ separation_{ij}}$ [a.u.] is the predicted separation between those same monomers in the CSynth conformation.

When comparing Hi-C contact maps (Supplementary Fig. S13 A, S13 G, S14), Relative Error = $\frac{2}{N(N+1)}\sum_{\mathrm{i,j \geq i}}^{\mathrm{N}} \frac{|\mathrm{Input\ CP_{ij}} - \mathrm{Output\ CP_{ij}}|}{\mathrm{Input\ CP_{ij}}} \times 100\%$. CP is contact probability and $\frac{2}{N(N+1)}\sum_{\mathrm{i,j \geq i}}^{\mathrm{N}}$ is an average over all unique, off diagonal, monomer pairs $(i, j) \in \{1, \ldots, N\}$ .

To compare CSynth conformations to absolute distances measured by FISH, we calculated Relative Error = $\frac{2}{N(N+1)}\sum_{\mathrm{i,j \geq i}}^{\mathrm{N}} \frac{|\mathrm{FISH\ separation_{ij}} - \mathrm{CSynth\ separation_{ij}}|}{\mathrm{FISH\ separation_{ij}}} \times 100\%$. We isotropically scaled CSynth conformations so their average pairwise separations match the average pairwise separations from FISH: $\frac{2}{N(N+1)}\sum_{\mathrm{i,j \geq i}}^{\mathrm{N}} \left|\mathrm{CSynth\ separation_{ij}}\right| = \frac{2}{N(N+1)}\sum_{\mathrm{i,j \geq i}}^{\mathrm{N}} \left|\mathrm{FISH\ separation_{ij}}\right|$. After this scaling, CSynth units match microscopy units (nanometers). When a FISH probe [bp] is located between CSynth monomers, its predicted CSynth location is inferred by interpolating between monomer locations using cubic splines.

**Human control Hi-C and FISH data**

Human IMR90 Hi-C data [S1] were accessed from the GEO using accession number: GSE63525. IMR90 FISH probe 3D locations [S2] were downloaded from: https://doi.org/10.5281/zenodo.3928890.

**Other eukaryotic Hi-C data**

Yellow fever mosquito, peanut, and roundworm Hi-C data were accessed from the GEO using accession number: GSE169088 [S3].

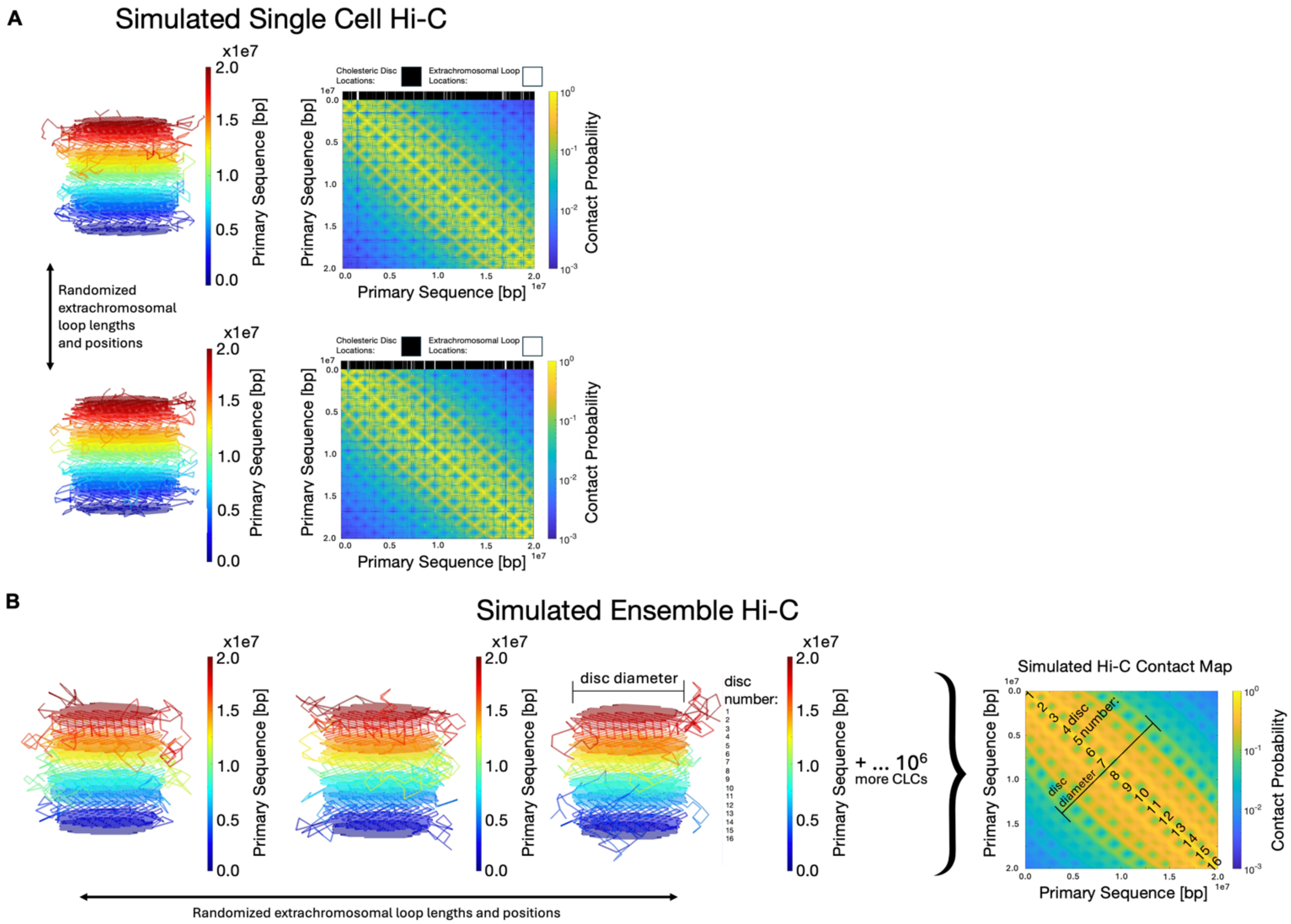


Figure S1. Simulated single cell versus population Hi-C for CLCs. A) Left: Example CLC structures with random extrachromosomal loop lengths and positions resulting in random cholesteric disc primary sequence locations. Right: Corresponding simulated single cell Hi-C contact maps. Top bar indicates location of extrachromosomal loops and cholesteric discs which vary across structures. B) Simulated Hi-C contact map for a population of $10^6$ CLC structures, approximating the number of sequenced cells in a typical bulk Hi-C experiment, each with random extrachromosomal loop lengths and positions and random cholesteric disc primary sequence locations. The cholesteric disc diameter and disc number are indicated on a structure and on the contact map.

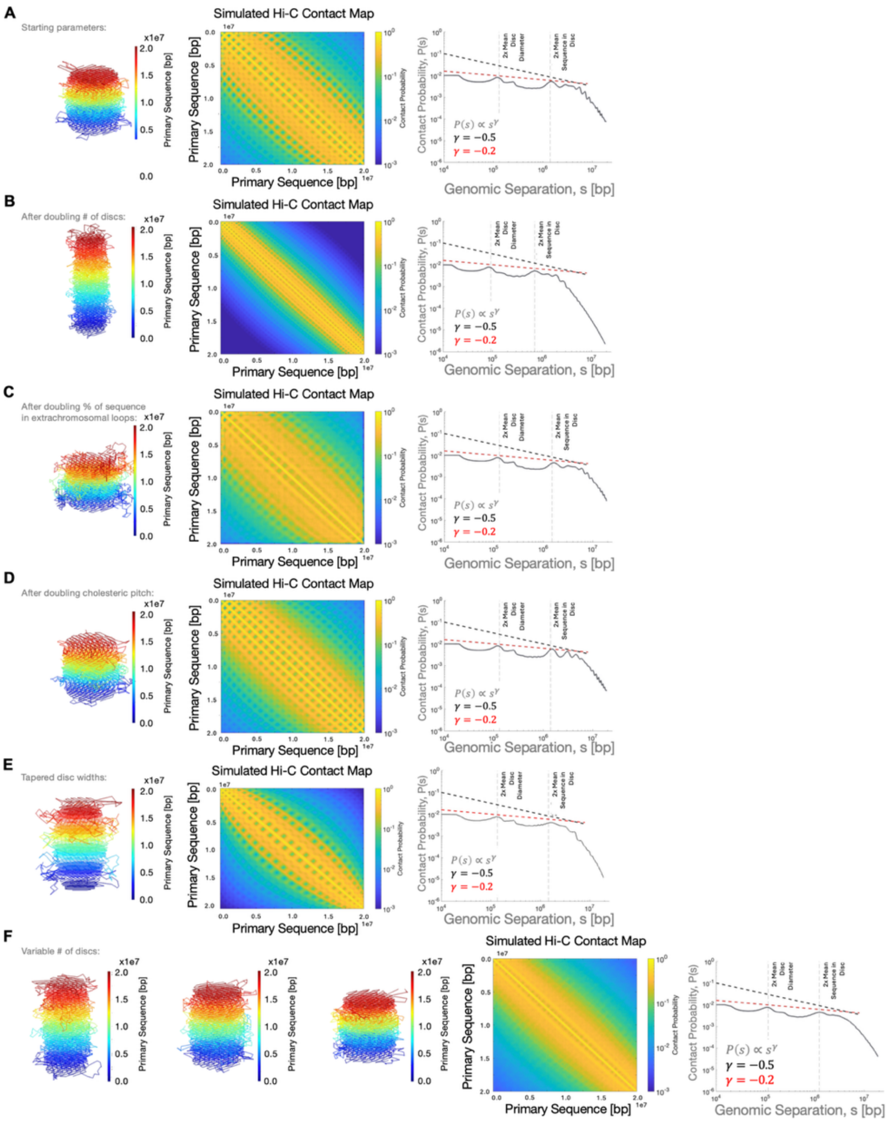


Figure S2. Many variants of the CLC model do not produce TADs. Contact probability scaling of CLC structures is robust to CLC parameters. Left: Example CLCs with random extrachromosomal loop lengths and positions. Middle: Simulated Hi-C data for a population ($N = 10^6$) of CLC structures. Right: Corresponding contact probability curve. Dinoflagellate contact probability curves obey a scaling exponent of $\gamma \approx -0.5$ (black dashed lines). Contact probability curves for a population of CLC structures obey a scaling exponent of $\gamma \approx -0.2$ (red dashed lines). The genomic separations corresponding to twice the average cholesteric disc diameter and to the average sequence in two cholesteric discs are indicated by vertical dashed lines. A) Starting parameters: total_chromosome_length = 4000 monomers, num_mon_per_disc = 150, frac_tot_sequence_in_loops = 0.1, frac_loop_sequence_inter_disc = 0.5, pitch = 0.65. B) After doubling the number of discs. C) After doubling the percentage of total sequence in extrachromosomal loops. D) After doubling the cholesteric pitch. E) After tapering disc widths at each chromosome end. F) For each CLC structure, the number of monomers per disc is a random number following a normal distribution with mean 150 monomers (corresponding to 27 discs) and standard deviation 40 monomers ($\pm$10 discs). As the total number of monomers is the equal across all CLC structures, a random number of monomers per disc across structures corresponds to a random number of cholesteric discs across structures.

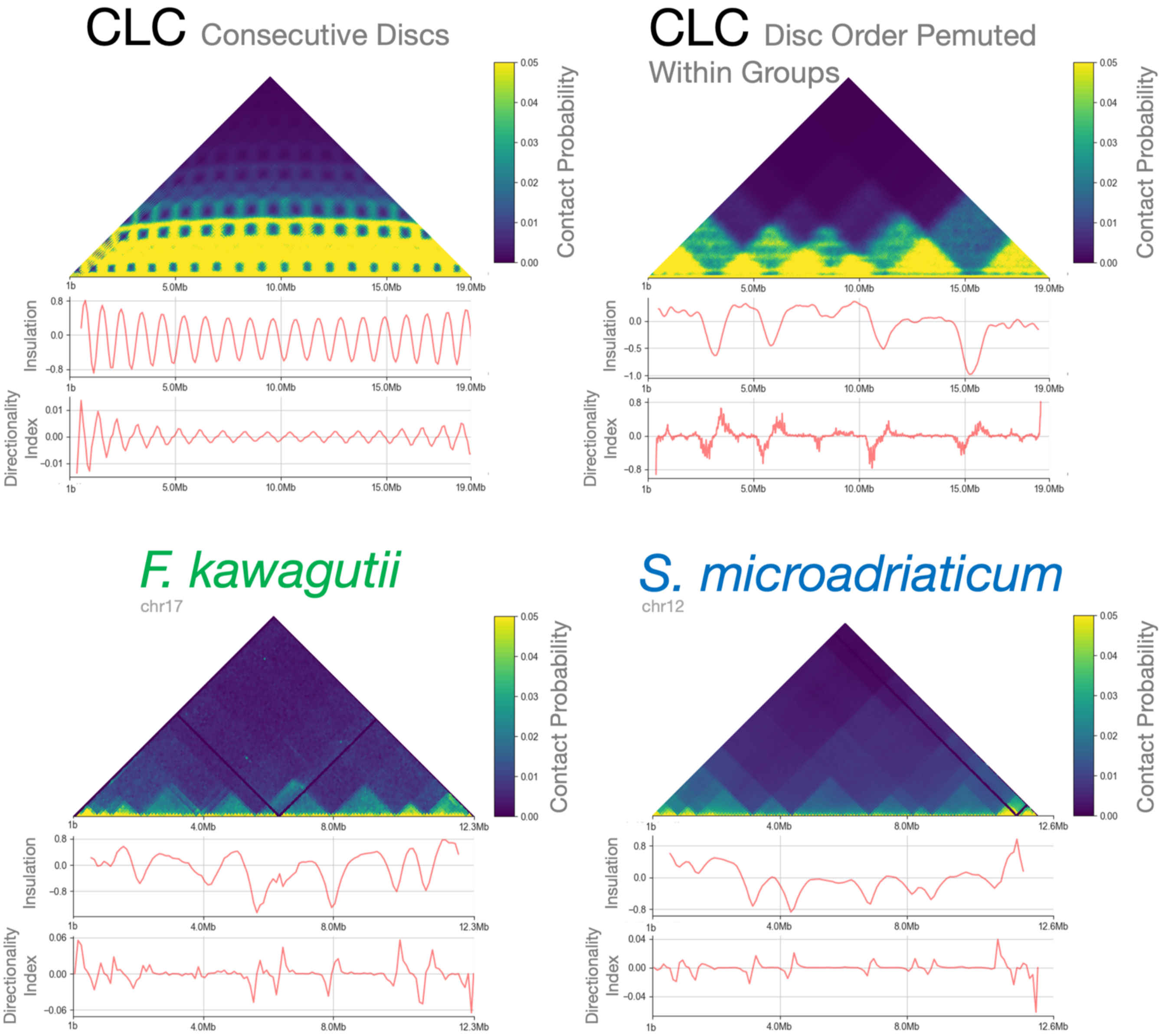


Figure S3. Insulation scores and directionally index from Hi-C contact maps for (top row) CLCs with consecutive discs or with disc order permuted within groups and (bottom row) *F. kawagutii* and *S. microadriaticum*. Black stripes in dinoflagellate contact maps indicate regions of unknown sequence.

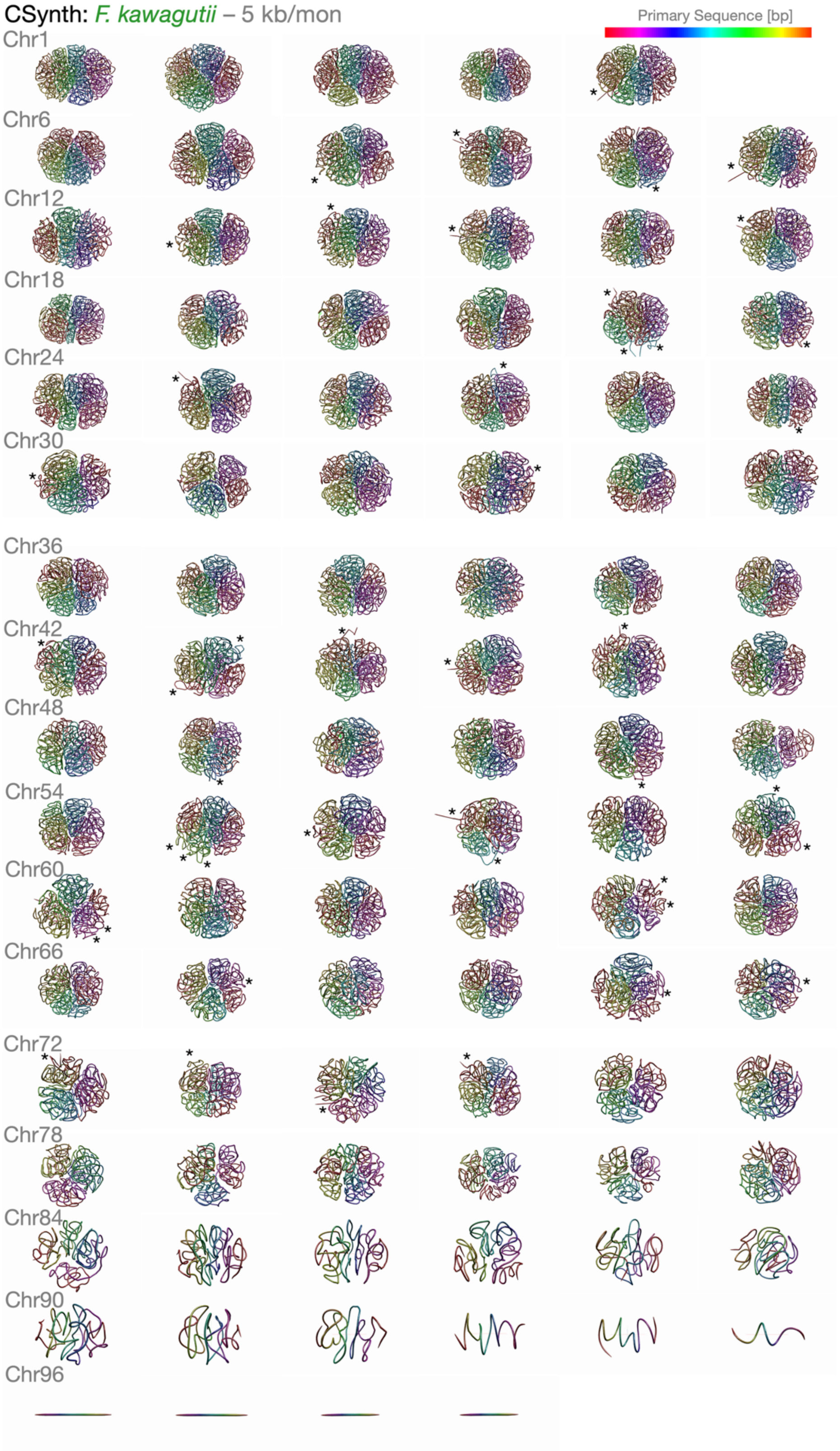


Figure S4. CSynth-generated conformations of *Fugacium kawaguti* chromosomes at 5 kb per monomer resolution. DNA is colored according to primary sequence. Very short Hi-C scaffolds (see Table S1) correspond to straight CSynth conformations (bottom row). Asterisks indicate extrachromosomal loops, observed in 35 of 99 chromosomes.

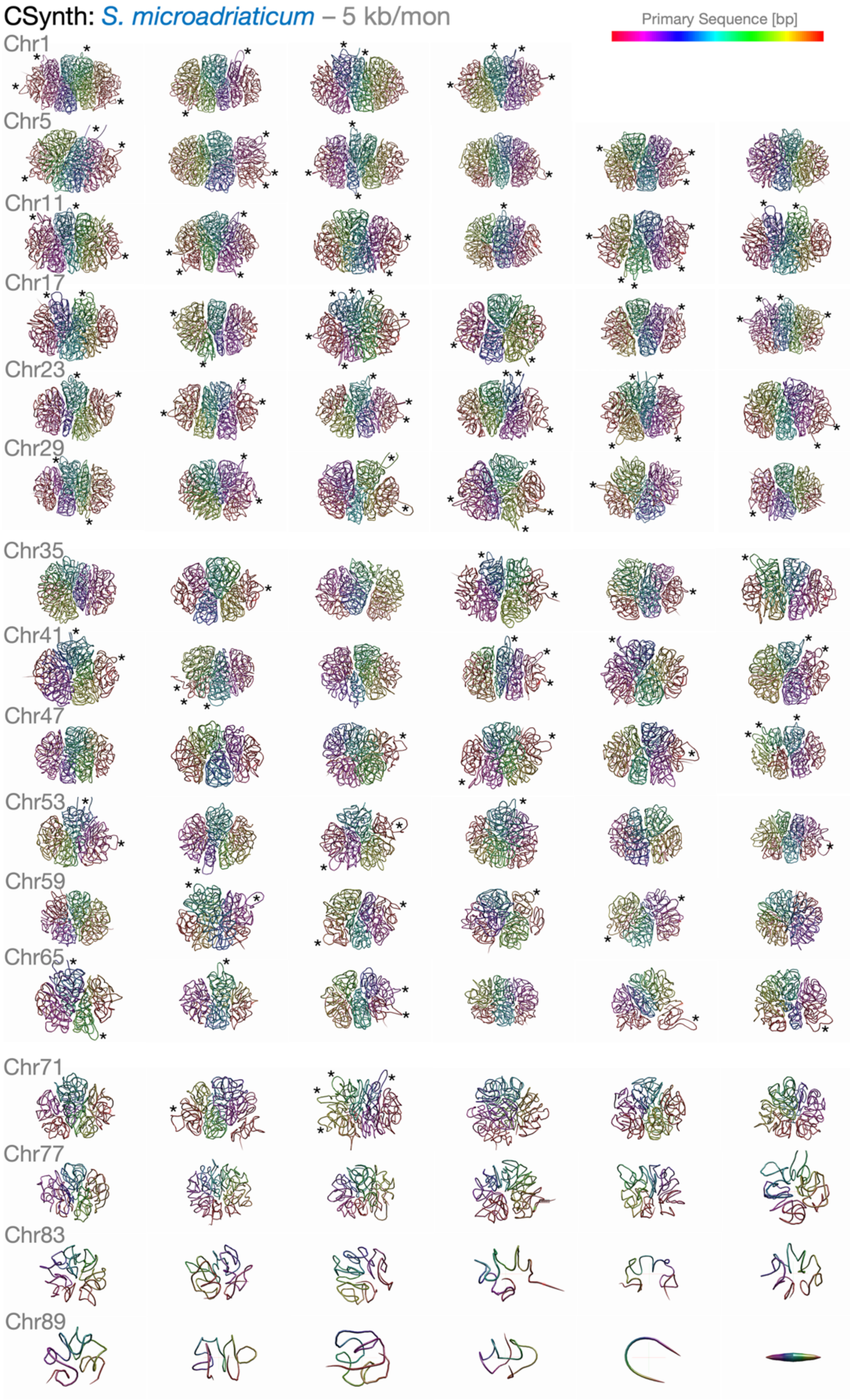


Figure S5. CSynth-generated conformations of *Symbiodinium microadriaticum* chromosomes at 5 kb per monomer resolution. DNA is colored according to primary sequence. Very short Hi-C scaffolds (see Table S2) correspond to straight CSynth conformations. Asterisks indicate extrachromosomal loops, observed in 62 out of 94 chromosomes.

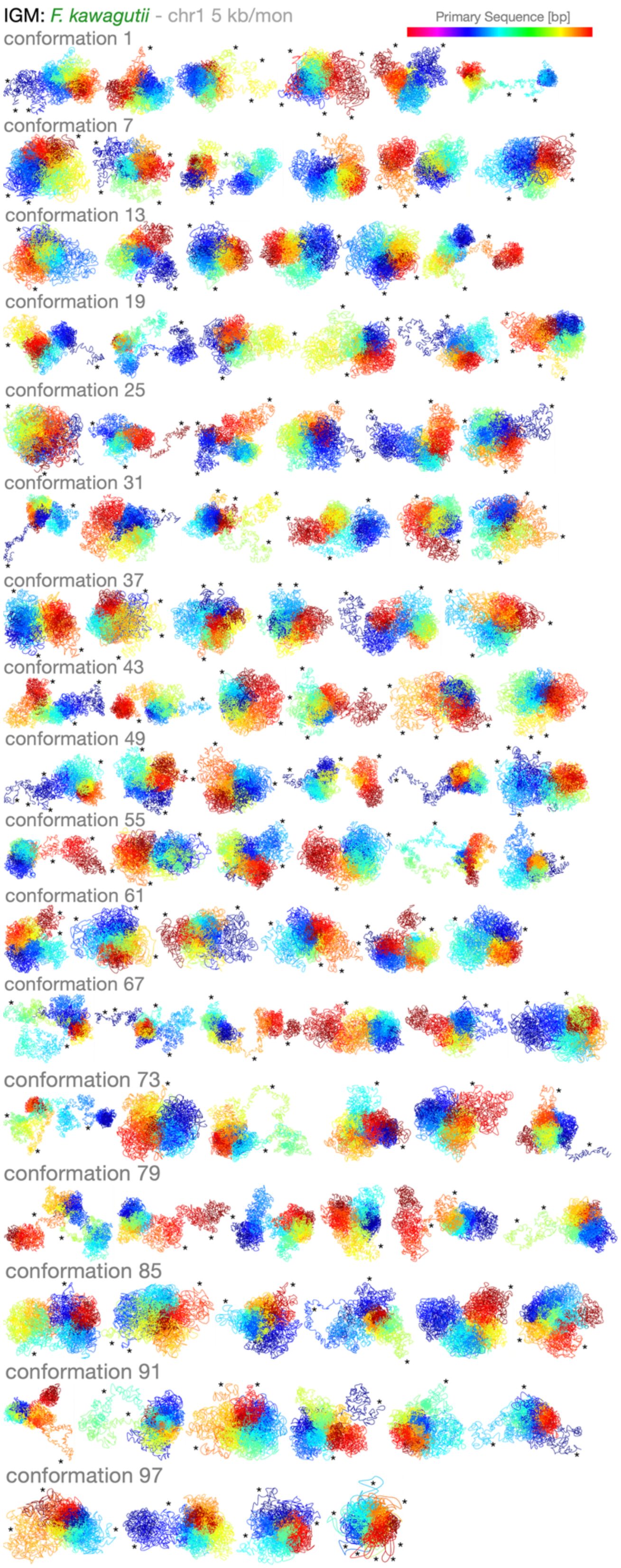


Figure S6. IGM-generated population (100 total conformations) of *Fugacium kawagutii* chromosome 1 at 5 kb per monomer resolution. DNA is colored according to primary sequence. Asterisks indicate extrachromosomal loops, observed in 100 of 100 conformations.

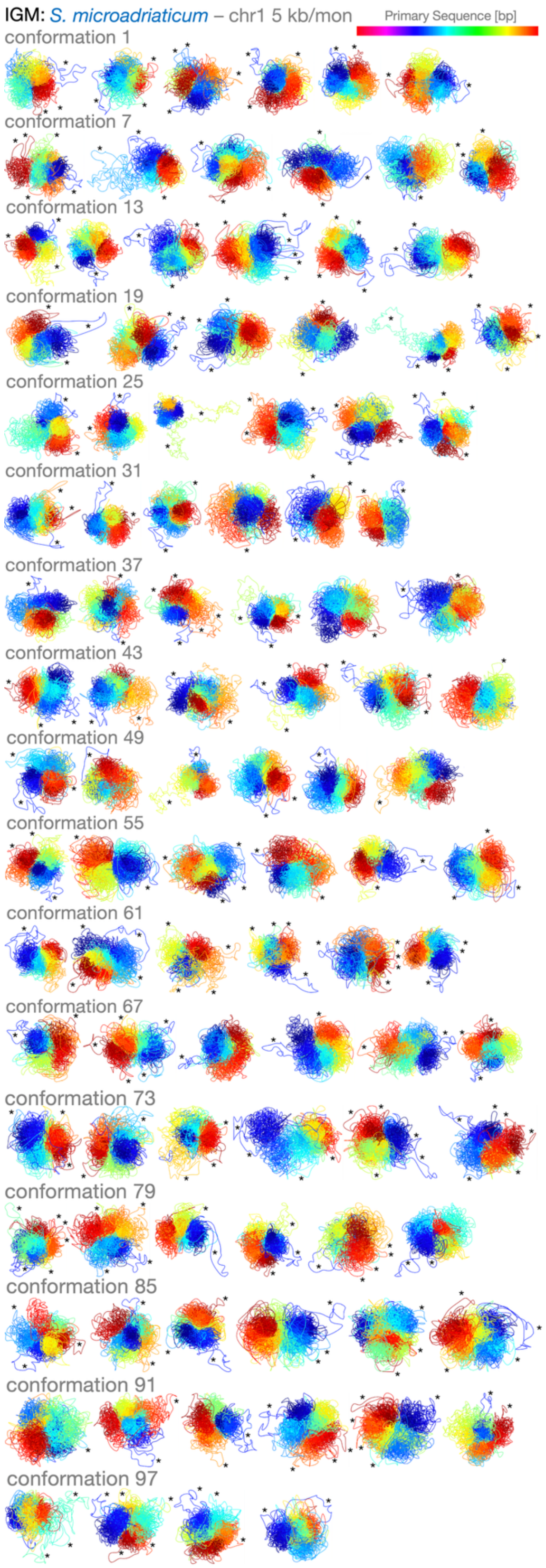


Figure S7. IGM-generated population (100 total conformations) of *Symbiodinium microadriaticum* chromosome 1 at 5 kb/monomer resolution. DNA is colored according to primary sequence. Asterisks indicate extrachromosomal loops, observed in 100 of 100 conformations.

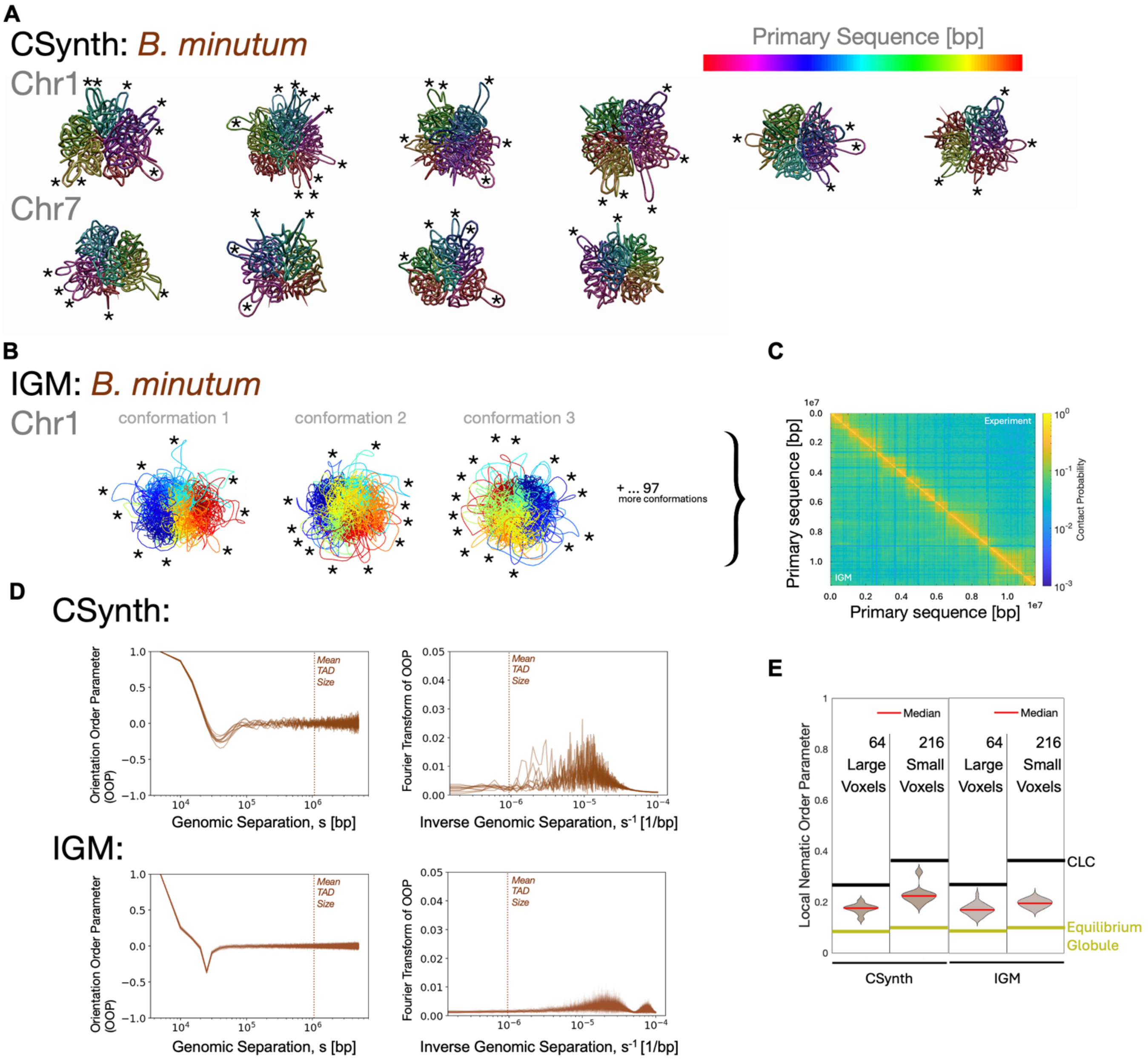


Figure S8. 3D reconstruction of large *Breviolum minutum* chromosome scaffolds. A) CSynth-generated conformations at 5 kb per monomer resolution. DNA is colored according to primary sequence. Asterisks indicate extrachromosomal loops. B) IGM-generated conformations (3 of 100 total) of *B. minutum* chromosome 1 at 5 kb per monomer resolution. C) Empirical Hi-C contact map of *B. minutum* chromosome (upper right triangle) and simulated Hi-C contact map from IGM populations (lower left triangles). Pearson correlation coefficient, $r = 0.92$. D) Orientational Order Parameter and its Fourier transform for CSynth and IGM conformations of *B. minutum* chromosome 1. E) Local nematic order parameter for CSynth and IGM conformations. Horizontal lines correspond to NOP for CLCs (black) and equilibrium globules (yellow).

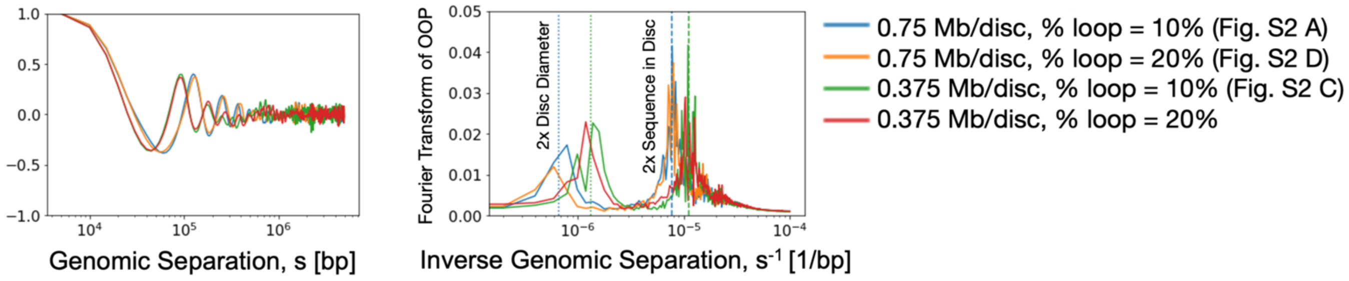


Figure S9. Orientation Order Parameter and its Fourier transform for CLC structures with varying parameters, namely the amount of sequence per disc and the percentage of total sequence in extrachromosomal loops. Dashed lines: reciprocal of the average amount of sequence in two cholesteric discs, reciprocal of the amount of sequence in twice the average disc diameter.

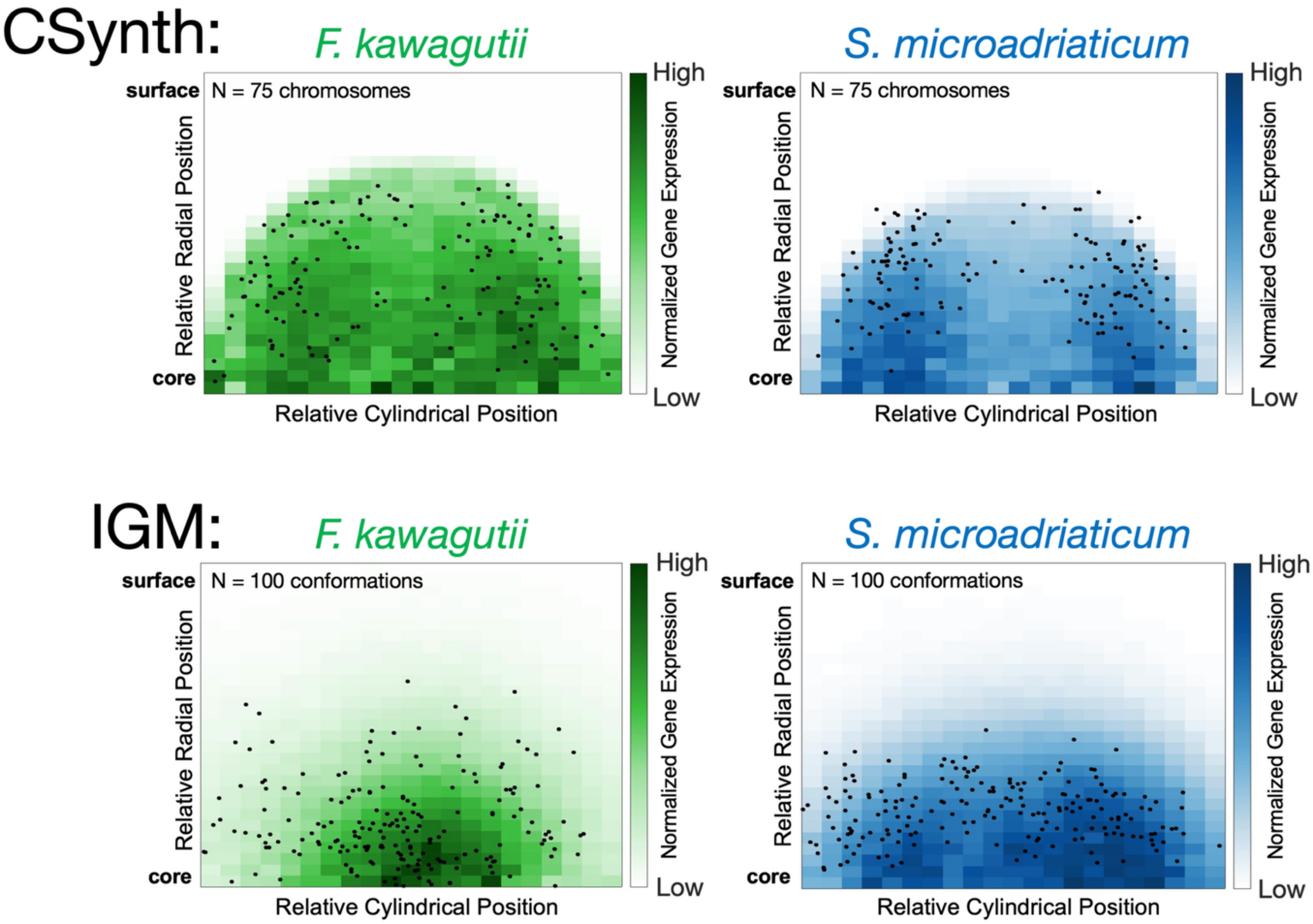


Figure S10. Telomere locations (black dots) superimposed onto active gene density plots from Figure 5 B and 5 C. Telomeres were defined as spanning 1% from each end of the chromosome which is on average 100 kb.

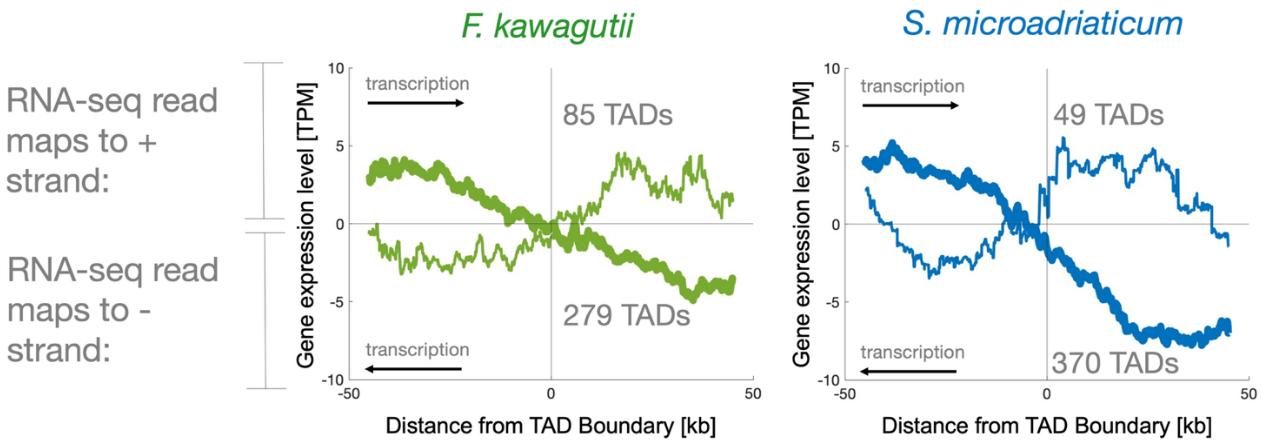


Figure S11. Using CSynth TAD annotations, we find TAD boundaries in *F. kawagutii* correspond to gene orientation switching at convergent gene arrays (thick lines) and divergent gene arrays (thin lines). This was found previously for *S. microadriaticum* [S7] and *B. minutum* [S8]. TAD boundaries at divergent gene arrays are less frequent than at convergent gene arrays (sample sizes indicated). The direction of transcription can be inferred from the strand to which RNA-seq reads are mapped and is given by the sign of the expression level (TPM).

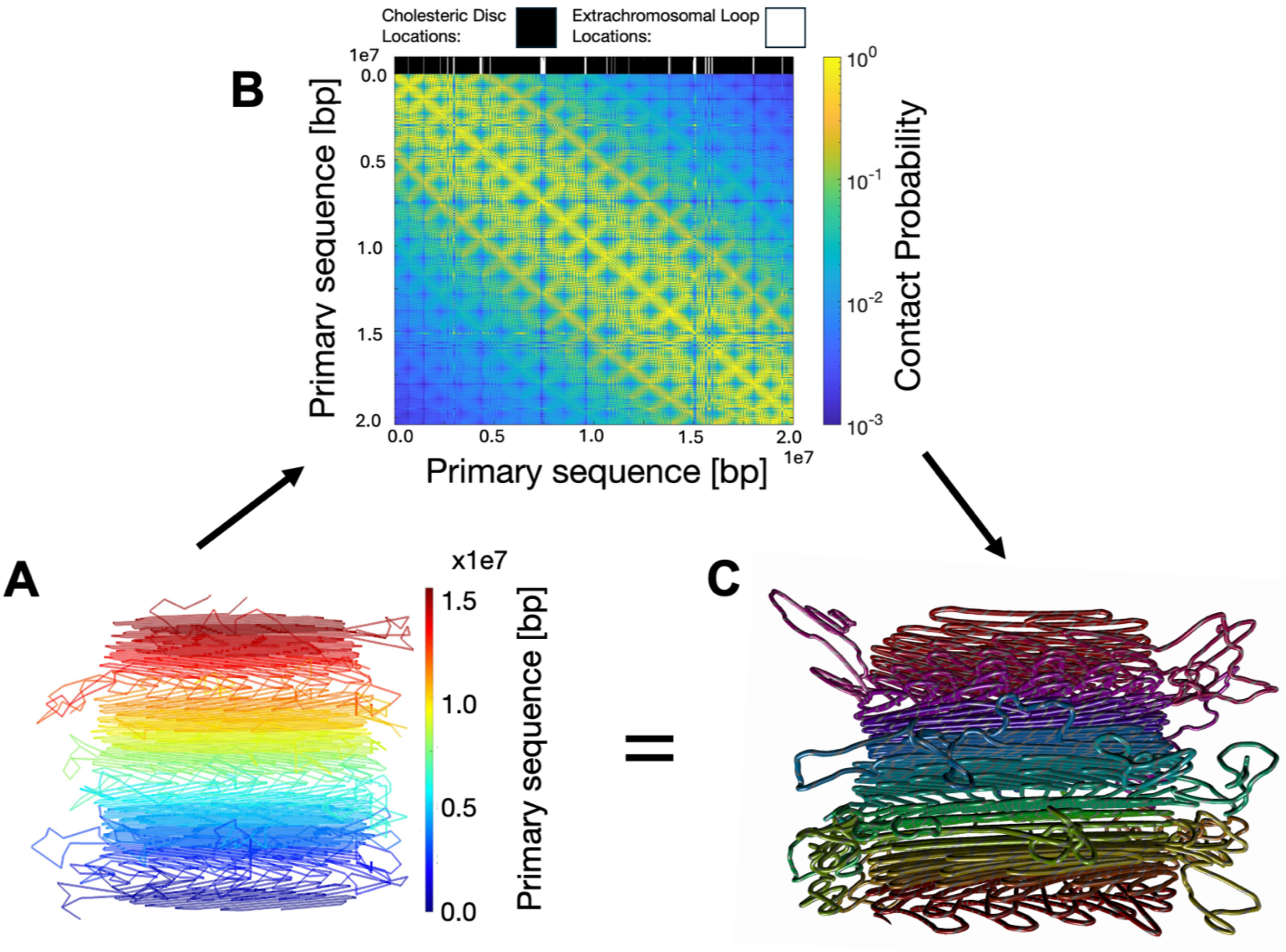


Figure S12. CSynth successfully reproduces input polymer conformation from simulated Hi-C contact map. A) CLC polymer structure. B) Simulated Hi-C contact map from the single CLC structure in A). Top bar indicates the primary sequence location of cholesteric discs and extrachromosomal loops. C) CSynth-predicted conformation using simulated Hi-C data in B). Here, monomer positions from A) are unknown to CSynth. The chromosome-averaged relative error between A) and C) is 7%.

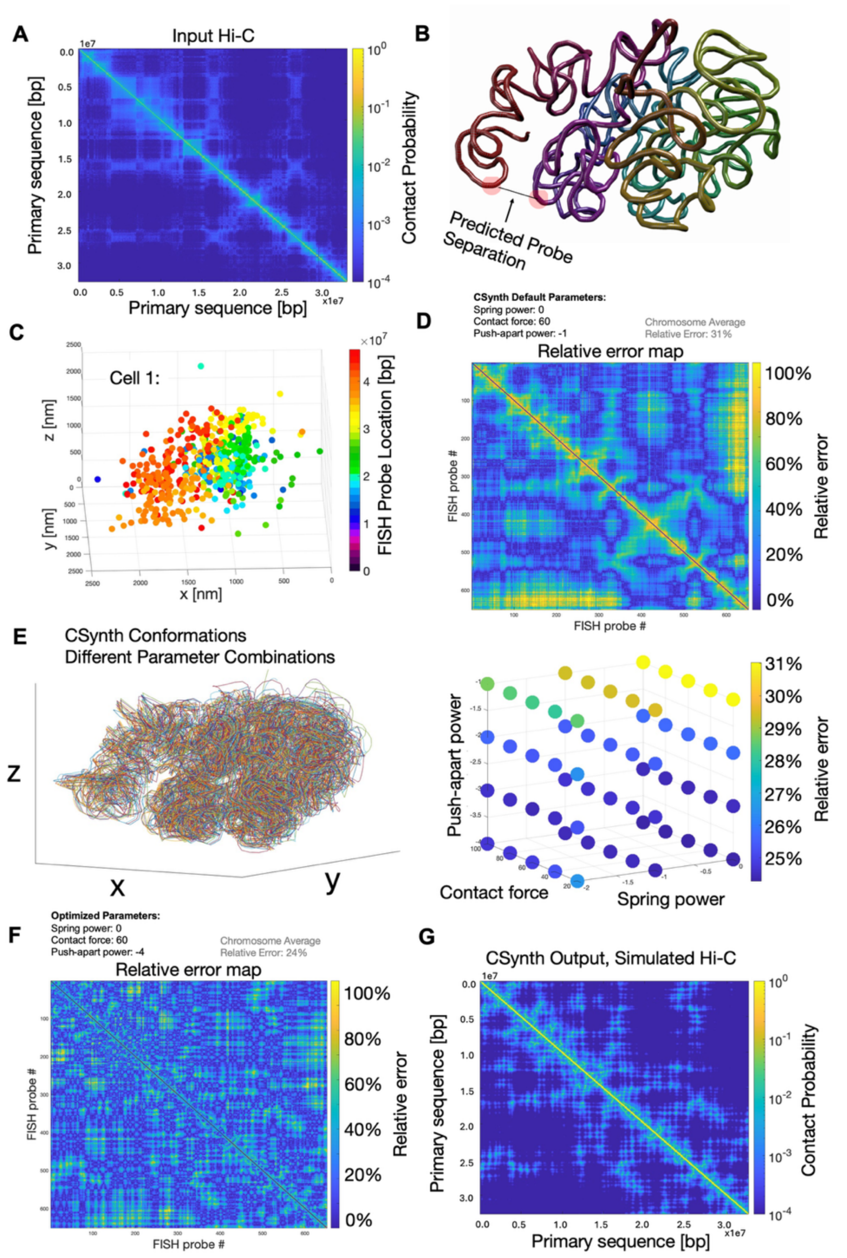


Figure S13. CSynth-predicted 3D chromosome conformations reproduce empirical FISH probe separations of 652 genomic loci on chr 21 in human cells. A) Hi-C data [S1] of chr 21 in human IMR90 cells were used as input to compute CSynth conformations. B) CSynth conformations make predictions for FISH probe separations that can be compared to cell-averaged separations of FISH probes. C) Cellular location of FISH probes in an example human IMR90 cell (chr 21) as measured by [S2]. D) The relative error map is shown for CSynth default parameters. E) CSynth conformations are sensitive to Push-apart power, but not other parameters. CSynth conformations (left) and chromosome-averaged relative errors (right) for different combinations of simulation parameters. Each ribbon, dot corresponds to a unique set of parameters. F) The relative error map for the optimal parameter set. G) Simulated Hi-C contact map for the CSynth conformation made using optimized parameters gives a contact map highly similar (Relative Error = 3%) to the input Hi-C in A).

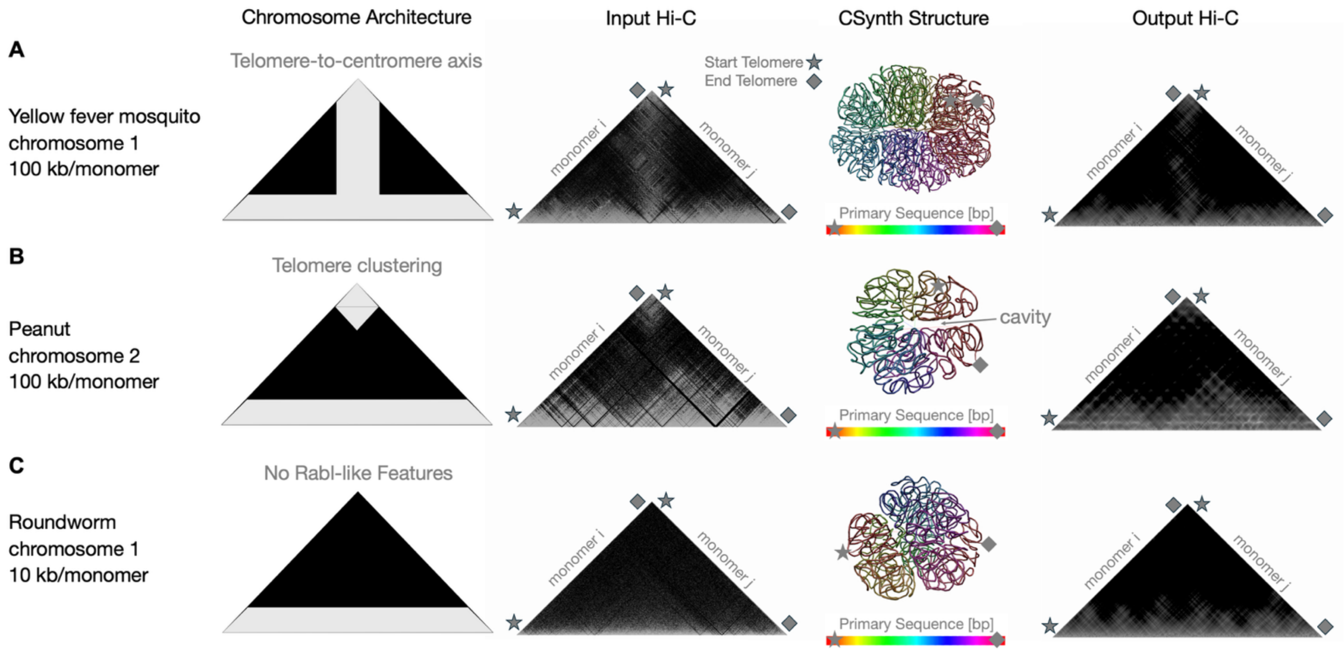


Figure S14. CSynth reproduces a wide variety of chromosome architectures from diverse eukaryotes. Far left: Distinct chromosome architectures. Left: Input Hi-C contact map (data: [S3]). Right: Corresponding CSynth conformations. Far right: Output Hi-C generated from CSynth conformation. Telomeres are indicated by the star and diamond shapes and CSynth conformations are colored according to primary sequence. A) Yellow fever mosquito chromosome 1 has high contact probabilities along the telomere-to-centromere axis. The predicted CSynth conformation (Relative Error = 10.6%) reflects this with its hairpin-shape and high density of contacts between opposite sides of the hairpin along the entire chromosome's length. B) Peanut chromosome 2 exhibits telomere clustering; in the predicted CSynth conformation (Relative Error = 8.3%), the hairpin allows telomeres to associate but there is a cavity (arrow) preventing other internal monomers from strongly interacting. C) Roundworm chromosome 1 exhibits no Rabl-like features (defined as centromere or telomere clustering) [S3]; the corresponding CSynth conformation (Relative Error = 11.4%) exhibits spatially segregated telomeres.

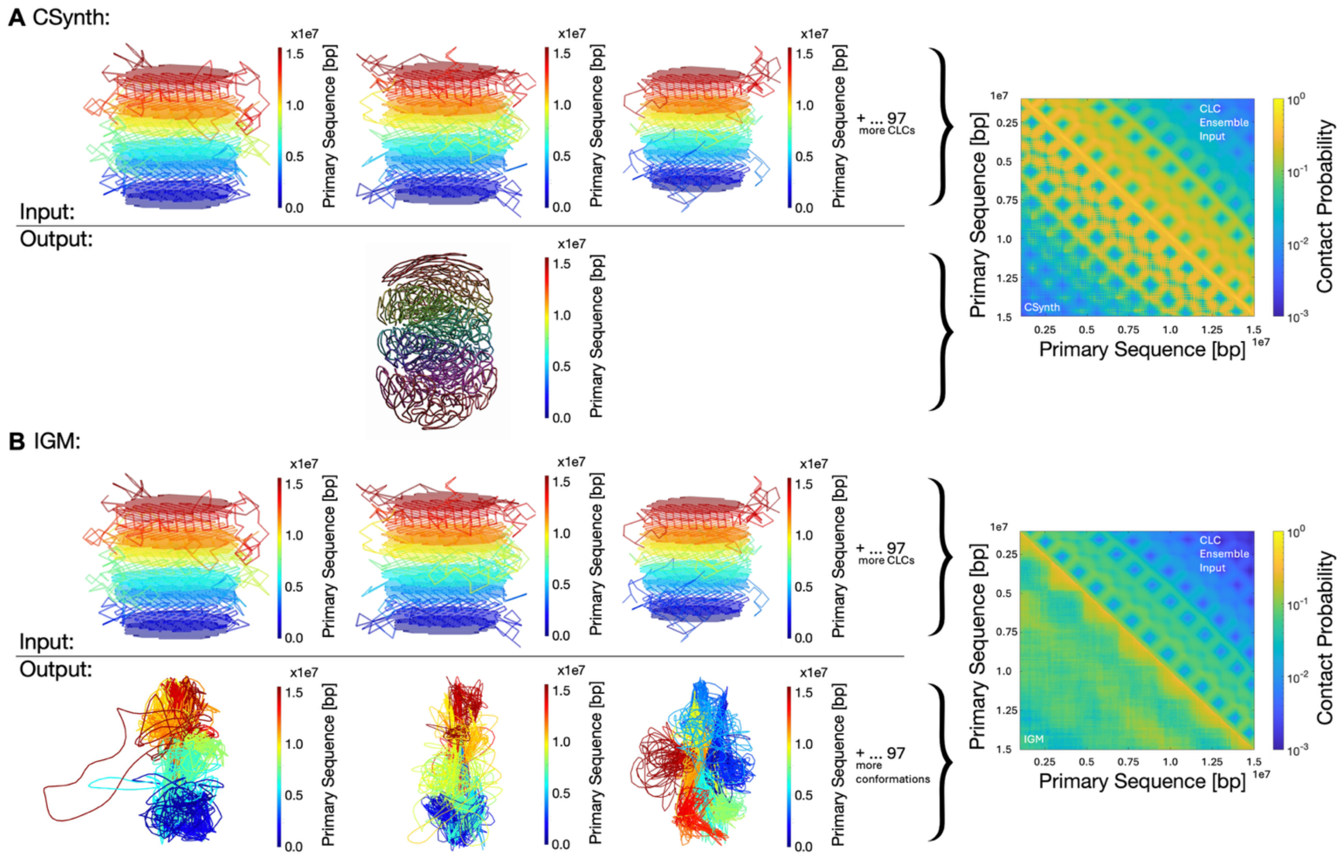


Figure S15. CSynth better reproduces a CLC structure from a simulated CLC population Hi-C contact map than IGM. A) CSynth prediction from CLC population input. Top: Input polymer conformations used to produce contact map input (top right triangle). Bottom: CSynth prediction from contact map input. Simulated Hi-C contact map from CSynth (bottom left triangle) correlates well with the CLC input contact map (Pearson correlation = 0.816). B) IGM prediction from CLC population input. Top: Input polymer conformations used to produce contact map input (top right triangle). Bottom: 3 select conformations from IGM population (100 total) generated from contact map input. Simulated Hi-C contact map from IGM (bottom left triangle) does not correlate with the CLC input contact map (Pearson correlation = 0.491).

**Table S1. Length of *F. kawagutii* Hi-C scaffolds.**

| Scaffold Number | Hi-C Scaffold Length [bp] | Scaffold Number | Hi-C Scaffold Length [bp] |
|---|---|---|---|
| 1 | 17515114 | 51 | 9873058 |
| 2 | 17193144 | 52 | 9666238 |
| 3 | 16516815 | 53 | 9656961 |
| 4 | 16193569 | 54 | 9540724 |
| 5 | 15438684 | 55 | 9499247 |
| 6 | 14342835 | 56 | 9275285 |
| 7 | 14325616 | 57 | 9046501 |
| 8 | 14189933 | 58 | 8877407 |
| 9 | 13932500 | 59 | 8763055 |
| 10 | 13807081 | 60 | 8713665 |
| 11 | 13453130 | 61 | 8608942 |
| 12 | 13267346 | 62 | 8551168 |
| 13 | 13240368 | 63 | 8524654 |
| 14 | 12740071 | 64 | 8138786 |
| 15 | 12710713 | 65 | 8010256 |
| 16 | 12619654 | 66 | 7967152 |
| 17 | 12329054 | 67 | 7758615 |
| 18 | 12230733 | 68 | 7666773 |
| 19 | 12228138 | 69 | 7623838 |
| 20 | 12229428 | 70 | 7510472 |
| 21 | 12206074 | 71 | 7143499 |
| 22 | 12097103 | 72 | 6917924 |
| 23 | 11981612 | 73 | 6916642 |
| 24 | 11874341 | 74 | 6827364 |
| 25 | 11805358 | 75 | 6767081 |
| 26 | 11806154 | 76 | 6020243 |
| 27 | 11729274 | 77 | 5995450 |
| 28 | 11704788 | 78 | 5613619 |
| 29 | 11575004 | 79 | 5326494 |
| 30 | 11431720 | 80 | 5332616 |
| 31 | 11258442 | 81 | 4972340 |
| 32 | 11202280 | 82 | 4614152 |
| 33 | 11089433 | 83 | 4366943 |
| 34 | 10907009 | 84 | 3301870 |
| 35 | 10891392 | 85 | 2746642 |
| 36 | 10755762 | 86 | 2518038 |
| 37 | 10739478 | 87 | 2497318 |
| 38 | 10725037 | 88 | 2348226 |
| 39 | 10665229 | 89 | 2111094 |
| 40 | 10609746 | 90 | 2020887 |
| 41 | 10541196 | 91 | 1497982 |

| | | | |
|---|---|---|---|
| 42 | 10499572 | 92 | 1427578 |
| 43 | 10410863 | 93 | 852295 |
| 44 | 10404246 | 94 | 736724 |
| 45 | 10395125 | 95 | 320241 |
| 46 | 10391202 | 96 | 137074 |
| 47 | 10369839 | 97 | 125000 |
| 48 | 10303574 | 98 | 125000 |
| 49 | 10217952 | 99 | 103000 |
| 50 | 10169143 | | |

**Table S2. Length of *S. microadriaticum* Hi-C scaffolds.**

| Scaffold Number | Hi-C Scaffold Length [bp] | Scaffold Number | Hi-C Scaffold Length [bp] |
|---|---|---|---|
| 1 | 19282064 | 51 | 7783240 |
| 2 | 14247422 | 52 | 7854610 |
| 3 | 13877534 | 53 | 7547234 |
| 4 | 14588374 | 54 | 7910744 |
| 5 | 14261437 | 55 | 7211228 |
| 6 | 13848878 | 56 | 7448770 |
| 7 | 13587423 | 57 | 7694657 |
| 8 | 13299794 | 58 | 7260962 |
| 9 | 12421019 | 59 | 7141439 |
| 10 | 12211192 | 60 | 6905598 |
| 11 | 12594407 | 61 | 6694038 |
| 12 | 12111265 | 62 | 6628099 |
| 13 | 11682917 | 63 | 6871460 |
| 14 | 11616559 | 64 | 6843148 |
| 15 | 11679755 | 65 | 6549511 |
| 16 | 11117062 | 66 | 6261677 |
| 17 | 11807614 | 67 | 6526534 |
| 18 | 11264987 | 68 | 6180782 |
| 19 | 10617761 | 69 | 5678934 |
| 20 | 10619008 | 70 | 6347547 |
| 21 | 10687167 | 71 | 5394230 |
| 22 | 10854226 | 72 | 5746528 |
| 23 | 10328379 | 73 | 5140143 |
| 24 | 10429750 | 74 | 4967601 |
| 25 | 10202498 | 75 | 4478657 |
| 26 | 10091992 | 76 | 4321980 |
| 27 | 10373796 | 77 | 3477828 |
| 28 | 9963233 | 78 | 3903991 |
| 29 | 10105166 | 79 | 3177586 |
| 30 | 10418649 | 80 | 2999361 |
| 31 | 9246414 | 81 | 2454569 |
| 32 | 10050580 | 82 | 1804558 |
| 33 | 9554063 | 83 | 1786823 |
| 34 | 9677557 | 84 | 1338160 |
| 35 | 9220381 | 85 | 1452270 |
| 36 | 8999273 | 86 | 1253655 |
| 37 | 9235022 | 87 | 1056224 |
| 38 | 8881437 | 88 | 929199 |
| 39 | 9335373 | 89 | 832037 |
| 40 | 8770219 | 90 | 782248 |

| | | | |
|---|---|---|---|
| 41 | 8858777 | 91 | 702567 |
| 42 | 8590938 | 92 | 521257 |
| 43 | 8845826 | 93 | 139507 |
| 44 | 8457715 | 94 | 27448 |
| 45 | 8421903 | | |
| 46 | 8221147 | | |
| 47 | 8672157 | | |
| 48 | 7644520 | | |
| 49 | 8347171 | | |
| 50 | 8442961 | | |

**Table S3. Estimated genomic coverage of Hi-C scaffolds.**

| Species | *F. kawagutii* | *S. microadriatica m* | *B. minutum* |
|---|---|---|---|
| k-mer genome size estimate [Gbp] [S9] | 1.05 | 1.18 | 1.50 |
| Genome fraction scaffolded by Hi-C | 0.79 | 0.67 | 0.39 |

**Table S4. Length of 75 largest *B. minutum* Hi-C scaffolds.**

| Scaffold Number | Hi-C Scaffold Length [bp] | Scaffold Number | Hi-C Scaffold Length [bp] |
|---|---|---|---|
| 1 | 11576343 | 41 | 7090168 |
| 2 | 11107596 | 42 | 7064639 |
| 3 | 11105505 | 43 | 6892915 |
| 4 | 10928834 | 44 | 6830015 |
| 5 | 10116126 | 45 | 6762807 |
| 6 | 9855553 | 46 | 6717247 |
| 7 | 9508929 | 47 | 6708150 |
| 8 | 9363653 | 48 | 6692000 |
| 9 | 9323510 | 49 | 6614245 |
| 10 | 9305992 | 50 | 6480833 |
| 11 | 9301534 | 51 | 6466334 |
| 12 | 9234120 | 52 | 6410878 |
| 13 | 8828819 | 53 | 6339702 |
| 14 | 8714911 | 54 | 6317251 |
| 15 | 8654588 | 55 | 6287587 |
| 16 | 8445662 | 56 | 6285956 |
| 17 | 8417553 | 57 | 6264918 |
| 18 | 8379267 | 58 | 5953329 |
| 19 | 8207709 | 59 | 5952774 |
| 20 | 8146391 | 60 | 5915379 |
| 21 | 8118824 | 61 | 5786850 |
| 22 | 8082874 | 62 | 5781716 |
| 23 | 8074619 | 63 | 5751648 |
| 24 | 7958149 | 64 | 5751630 |
| 25 | 7879853 | 65 | 5748650 |
| 26 | 7809123 | 66 | 5645890 |
| 27 | 7805800 | 67 | 5590159 |
| 28 | 7721756 | 68 | 5573185 |
| 29 | 7706142 | 69 | 4804405 |
| 30 | 7699351 | 70 | 4668170 |
| 31 | 7614309 | 71 | 4647415 |
| 32 | 7585209 | 72 | 4577699 |
| 33 | 7509923 | 73 | 4563830 |
| 34 | 7474210 | 74 | 4490779 |
| 35 | 7455639 | 75 | 4299974 |
| 36 | 7453953 | | 4109746 |
| 37 | 7207301 | | |
| 38 | 7134339 | | |
| 39 | 7134015 | | |
| 40 | 7102662 | | |
